# Immunotherapy restores therapeutic efficacy of fecal microbiome transplants to treat *Clostridioides difficile*

**DOI:** 10.64898/2026.09.17.752434

**Authors:** Subham Mridha, Md Zahidul Alam, Joshua E. Denny, Ellie N. Hulit, Jeffrey R. Maslanka, Nontokozo V. Mdluli, Minh Tran-Ha, Kevin S Mears, Daivon D. Brown, Mohamad-Gabriel Alameh, Michael C. Abt

**Author notes:** Corresponding Author: Department of Microbiology, Perelman School of Medicine, University of Pennsylvania, Philadelphia, Pennsylvania, USA., 303B Johnson Pavilion, 3610 Hamilton Walk. Philadelphia, Pennsylvania 19104. Author contributed equally to work (Subham Mridha, Md Zahidul Alam).

## Abstract

Microbiome-based therapeutics to treat enteric bacterial infections requires stable engraftment of transplanted beneficial bacteria that target and eliminate the pathogen. The host factors that determine whether a microbiome transplant engrafts remain largely unresolved. Here we demonstrate that Interleukin-10 (IL-10) signaling deficiency leads to impaired fecal microbiota transplantation (FMT) engraftment and failure to resolve *Clostridioides difficile* infection in mice. In the absence of IL-10-mediated immunoregulation provided by Foxp3^+^ T_reg_ cells, increased IFN-γ signaling in the intestine leads to elevated production of reactive oxygen/nitrogen species (ROS/RNS) by neutrophils and epithelial cells that supports the growth of inflammation-tolerant microbes, inhibits FMT engraftment, and impairs resolution of *C. difficile.* FMT treatment success can be restored by antibody-mediated blockade of IFN-γ signaling, neutrophil depletion or inhibition of ROS/RNS production. Lastly, we developed an *Il10*-mRNA-LNP immunotherapy to boost IL-10 signaling in FMT Non-Responsive mice and demonstrate that *Il10*-mRNA-LNP administration is sufficient to restore FMT engraftment and subsequent resolution of *C. difficile*. Combined, these data support a mechanism by which IL-10 released by T_reg_ cells limits IFN-γ mediated intestinal inflammation thereby promoting an intestinal microenvironment receptive to FMT engraftment and resolution of *C. difficile*. These data demonstrate that the host’s immune status can be therapeutically modulated to improve microbiome-based approaches to treat infection.

## Introduction

Microbiome-based therapeutics, which encompass fecal microbiota transplant (FMT) and defined live biotherapeutic products, have the potential to treat a wide range of chronic diseases associated with intestinal dysbiosis by resetting the diseased individual’s microbiome^1,2^. However, to date, such microbiome-based approaches have been approved to treat only a narrow spectrum of these diseases, specifically recurrent *Clostridioides difficile* infection^3–8^. In past 4 years, the U.S. Food and Drug Administration has approved two standardized fecal microbiota–based therapies for the prevention of *C. difficile* recurrence^9–11^. Clinical trials to test the efficacy of microbiome-based therapeutics to treat other diseases have had limited success^12^. Therefore, the underlying biological mechanisms that determine the efficacy of microbiome therapies need to be better understood. While advances have been made in identifying beneficial microbial consortia^11,13–25^, host factors that shape the success of these therapies remain poorly defined. The immune system, particularly the balance between pro and anti-inflammatory pathways, influences whether transplanted microbes engraft and exert their beneficial effects^1,26^. Understanding the immunological mechanisms that determine microbiome treatment outcomes is crucial to develop predictable, safe, and effective interventions for dysbiosis-associated diseases.

Inflammatory Bowel Disease (IBD) patients experience up to eightfold higher rates of *C. difficile*-related complications, which include more severe disease, prolonged hospitalizations, and increased risk of *C. difficile* recurrence^27–31^. These worse outcomes can occur even without prior antibiotic exposure, likely due to gut barrier disruption, heightened inflammation, and alterations in the intestinal microbiome^32,33^. While FMTs achieve high cure rates in patients with uncomplicated, recurrent <EM>C. difficile</EM>^34^, its application in patients with underlying IBD remains limited due to safety concerns and has yielded inconsistent outcomes in the few trials in which IBD patients were included^35–38^. As a result, microbiome-based therapeutics are often not a treatment option for IBD patients experiencing an episode of *C. difficile*. In the absence of a microbiome-based therapy, treatment options for IBD patients with *C. difficile* are limited and suboptimal. Standard antibiotic regimens used to treat *C. difficile* may further exacerbate dysbiosis in IBD^39^ and Bezlotoxumab, a monoclonal anti-toxin antibody therapy that reduces recurrence risk^40–42^, was discontinued from clinical use as of January 2025^43^. This lack of treatment modalities highlight the need to identify new microbiome-based therapeutic strategies that account for the host’s immune status to protect IBD patients from *C. difficile* infection.

In *C. difficile* infected mice, FMT transplant “Responders” or “Non-Responders” can be differentiated based on their pre-FMT immune status with our group reporting that Foxp3^+^ T-regulatory (T_reg_) cells support the success of an FMT in the context of *C. difficile* infection^26^. In this study we use the IL-10 deficient (*Il10*^-/-^) mouse IBD model and demonstrate that IL-10 signaling is required to support FMT engraftment and resolve *C. difficile* infection. Despite receiving identical FMT therapy, mice can shift between FMT Responders and Non-Responders by targeting the IL-10-IFNγ signaling axis prior to FMT treatment. Further, we develop an *Il10-*mRNA-lipid nanoparticle (LNP) immunotherapy that can convert FMT Non-Responders into Responders, demonstrating that therapeutically targeting immune pathways can improve efficacy of microbiome-based therapeutics.

## Results

### IBD driven loss of immune regulation renders *C. difficile* infected host non-responsive to FMT therapy

Current FMT guidelines to treat *C. difficile* infection often exclude patients with IBD^35^ and clinical trials suggest FMT in IBD patients have decreased host engraftment efficiency and decreased efficacy at preventing recurrence^36–38^. Genetic ablation of the immunosuppressive cytokine Interleukin-10 (IL-10) in mice leads to microbiome dependent intestinal inflammation that share many of the clinical features of IBD and has been extensively used to study aspects of IBD^44–47^. Therefore, we tested whether *C. difficile* infected *Il10^-/-^*mice exhibited an impaired capacity to respond to FMT therapy similar to IBD patients. *Il10* deficient (*Il10*^-/-^) and littermate control *Il10* heterozygous (*Il10*^HET^) mice were treated with an antibiotic regimen to disrupt the microbiome, then infected with *C. difficile* (CD196) spores via oral gavage (**Suppl Fig 1A**). At day 21 post-infection (p.i.), following establishment of persistent *C. difficile* colonization in the large intestine (**Suppl. Fig 1B**), mice were administered an FMT. *Il10*^HET^ and *Il10*^-/-^mice were cohoused prior to and throughout infection until being separated into individually housed cages on the day of FMT. *C. difficile* burden and intestinal microbiome composition were monitored in the feces to determine success of FMT engraftment and resolution of infection. *Il10*^HET^ mice resolved *C. difficile* burden below the limit of detection (**Fig. 1A**), similar to previous reports in both wild type mice and humans^4–7,15,26,34,48^. In contrast, *Il10*^-/-^mice maintained high levels of *C. difficile* burden following FMT demonstrating that the lack of IL-10 leads to FMT failure (**Fig. 1A**). Microbial engraftment of a bacterial transplant inoculum is critical for FMT treatment success^48–51^, therefore 16S rRNA sequencing of feces prior to FMT (Day 0 post FMT) and Day 8 post-FMT from FMT recipient mice was conducted to determine FMT engraftment. Prior to FMT, *C. difficile* infected *Il10*^HET^ and *Il10*^-/-^mice exhibited distinct microbiome profiles from each other and the FMT input (**Fig. 1B, Suppl. Table 1 PERMANOVA**). Post FMT, the intestinal microbial community of *Il10*^HET^ mice significantly shifted to resemble the composition of the FMT inoculum indicating successful engraftment while microbial community of *Il10*^-/-^mice remained distinct (**Fig 1B, C**). Unweighted UniFrac principal coordinate analysis (**Suppl. Fig. 1C**) and unweighted UniFrac distance relative to the FMT inoculum (**Suppl. Fig. 1D**) confirmed successful FMT engraftment in *Il10*^HET^ but not *Il10*^-/-^mice. Notably, ABX-treated, uninfected *Il10*^-/-^mice were capable of engrafting an FMT, although at a delayed rate compared to ABX-treated, uninfected *Il10*^HET^ mice suggesting that IL-10 genetic deficiency in combination with an ongoing *C. difficile* infection is needed to drive FMT failure (**Suppl. Fig 1E, F**).

**Fig1:**
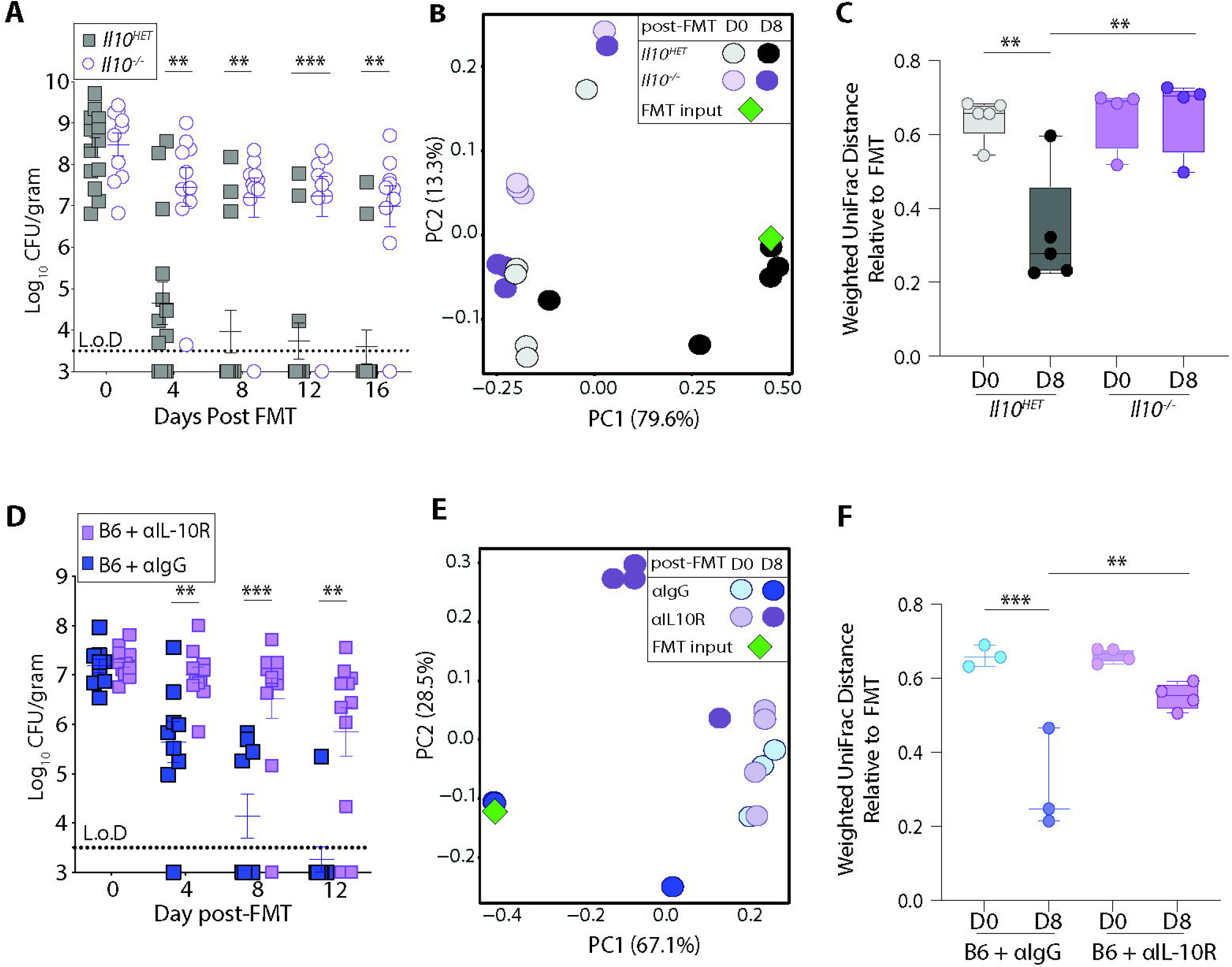
Loss of Interleukin 10 (IL-10) signalling leads to failed FMT engraftment and resolution of *C. difficile* infection. **(A)** *C. difficile* burden in fecal pellets from *Il10^HET^* (n=14) and *Il10^-/-^*(n=10) mice post FMT. Data is combined from two independent experiments. Experimental design shown in Supplemental **Fig S1A**. **(B)** Weighted UniFrac principal coordinate analysis plot of 16S bacterial rRNA ASVs from fecal pellets of *Il10^HET^* (n=5) and *Il10^-/-^*(n=4) mice at Day 0 and 8 post-FMT. **(C)** Weighted Unifrac distance comparing the microbial beta diversity dissimilarity of *Il10^HET^* and *Il10^-/-^*groups to the FMT inoculum. **(D)** *C. difficile* burden post FMT, in fecal pellets from C57BL/6 mice treated with αIgG (n=9) and αIL-10R (n=11) antibodies. Data is combined from 3 independent experiments. Experimental design shown in Supplemental **Fig S1I**. **(E)** Weighted UniFrac principal coordinate analysis plot of 16S bacterial rRNA ASVs from fecal pellets of C57BL/6 mice treated with αIgG (n=3) and αIL-10R (n=4) antibodies at Day 0 and 8 post-FMT. **(F)** Weighted Unifrac distance comparing the microbial beta diversity dissimilarity of αIgG and αIL-10R groups to the FMT inoculum. **(A, D)** Data are presented as mean values ± standard error of mean (SEM). *C. difficile* burden statistical significance was calculated by two-part model *t*-test. UniFrac distance statistical significance was calculated by one-way ANOVA test. * p < 0.05, **p <0.01, *** p < 0.001. **(C, F)** Boxes represent median and the first and third quartile. Whiskers extend to the highest and lowest data point. L.o.D.-Limit of Detection.

Germline *Il10* deficiency causes defects in intestinal immune homeostasis and drives microbiome dysbiosis immediately following weaning^44^. This pre-existing inflammation and microbial dysbiosis may confound interpretation of the contribution of IL-10 during FMT mediated clearance of *C. difficile*. To address this concern, wild type C57BL/6 mice were infected with *C. difficile* and following establishment of infection were administered anti-IL-10R or isotype (Rat IgG) antibody treatment to block IL-10 signaling after acute disease but prior to FMT. These mice were littermates and cohoused throughout the course of infection until FMT, to ensure the same starting microbiome (**Suppl. Table 1, PERMANOVA**). Prior to FMT, two weeks of IL-10R blockade did not alter *C. difficile* burden or microbiome composition (**Fig. 1D, E**). Following FMT, C57BL/6 mice treated with isotype control antibody resolved *C. difficile* (**Fig. 1D**) and successfully engrafted the FMT input (**Fig. 1E**, **F**). In contrast C57BL/6 mice treated with anti-IL10R antibody starting at day 7 post-infection and then administered an FMT failed to resolve *C. difficile* infection and their microbiome remained distinct from the FMT input (**Fig. 1D-F**). Unweighted UniFrac principal coordinate analysis (**Suppl. Fig. 1G**) and unweighted UniFrac distance relative to the FMT inoculum (**Suppl. Fig. 1H**) all supported the conclusion that anti-IL-10R treatment impaired FMT engraftment. Also, IL-10R blockade initiated on the day of FMT did not impair FMT mediated resolution of *C. difficile* infection (**Suppl. Fig 1I, J**). Combined these data demonstrate that a host can be rendered non-responsive to identical FMT inoculums by blocking a key immune pathway, and that IL-10 signaling prior to FMT administration is necessary to establish an intestinal environment that is receptive to FMT engraftment, which promotes resolution of *C. difficile* infection.

### T_reg_ cells are the main source of IL-10 following *C. difficile* infection

IL-10 is an immunoregulatory cytokine expressed by cells of both adaptive and innate immune system^45^. However, the relative proportion of IL-10 competent immune cell populations in the large intestine following *C. difficile* infection and prior to FMT is unknown. To determine the cellular source of IL-10 critical for FMT success, we quantified IL-10 producing cells in the large intestine lamina propria (LI Lp) following *C. difficile* infection using *IL10^GFP^* reporter mice. On the day of FMT (day 21 p.i.), the predominant IL-10-GFP^+^ cell population was CD4^+^ Foxp3^+^ T-regulatory (T_reg_) cells (**Fig 2A**, **Suppl Fig 2A**). This population of IL-10-GFP^+^ Foxp3^+^ T_reg_ cells significantly expanded in frequency (**Fig. 2B**) and total numbers (**Fig. 2C**) following infection while other IL-10-GFP^+^ immune cell populations remained relatively stable (**Suppl. Fig 2B**). Phenotypic characterization of IL-10-GFP^+^ Foxp3^+^ T_reg_ cells identified a highly immunosuppressive cellular phenotype at the day of FMT. IL-10-GFP^+^ Foxp3^+^ T_reg_ cells had elevated expression of transcription factors Rorγt and T-bet compared to IL-10-GFP^Neg^ Foxp3^+^ T_reg_ cells isolated from the same *C. difficile* infected mice, demonstrating that the IL-10 population was enriched in two subsets of peripherally induced T_reg_ cells that are important for regulating intestinal inflammation following infection^52–55^. Further comparison revealed that IL-10-GFP^+^ T_reg_ cells also had elevated expression of ICOS, CTLA-4 and lower expression of PD-1 (**Fig. 2D**), a co-inhibitory expression profile that correlates with enhanced immunosuppressive activity^56–63^. IL-10-GFP^+^ T_reg_ cells also significantly increased expression of co-inhibitory receptors CTLA-4 and ICOS following infection compared to IL-10 GFP^+^ T_reg_ cells isolated from uninfected mice (**Suppl. Fig. 2C**). In summary, phenotypic characterization of IL-10-GFP^+^ T_reg_ cells following *C. difficile* infection identified T_reg_ cells with an expression profile associated with enhanced immunosuppressive activity.

**Fig2:**
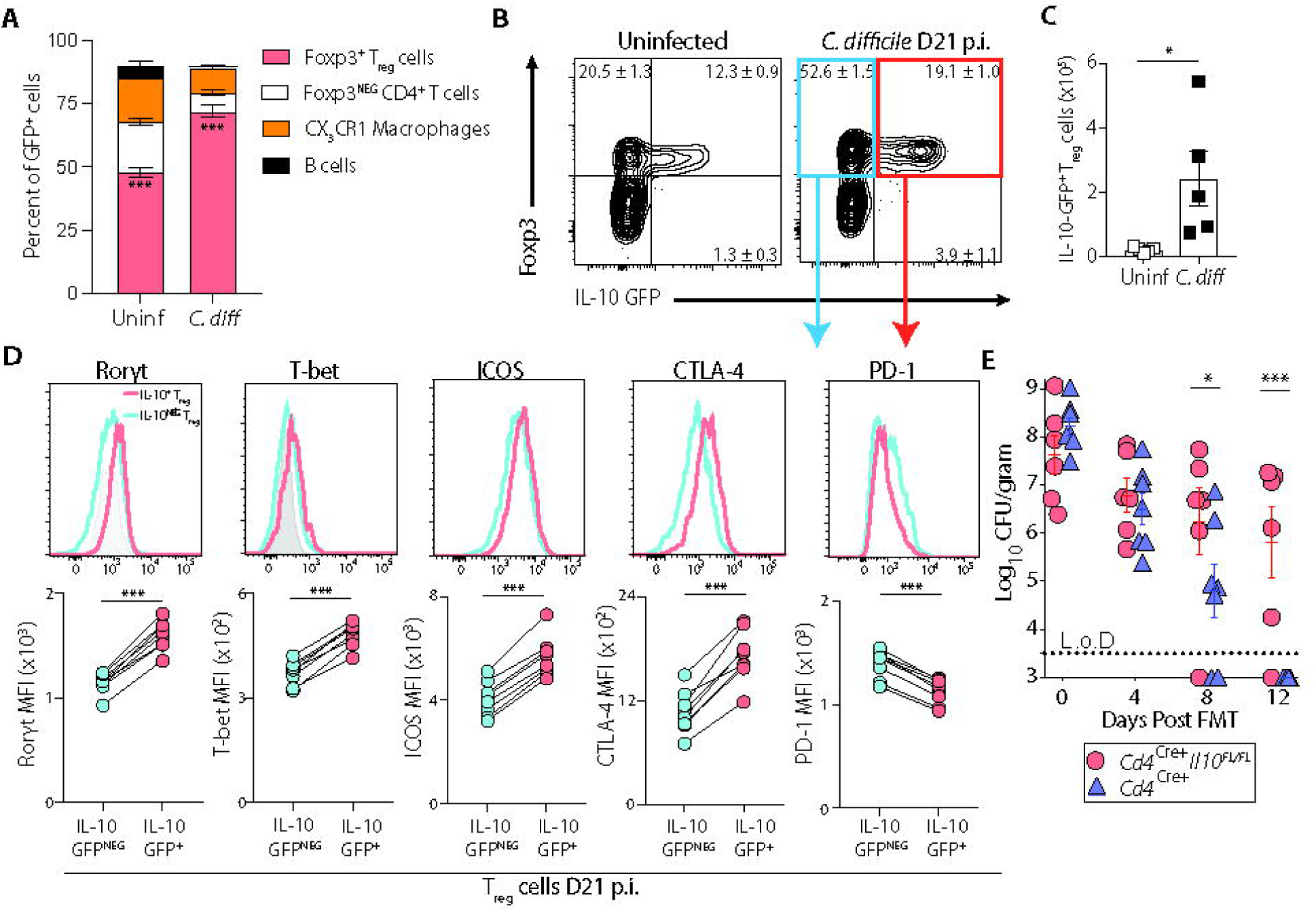
Treg cells are the main source of IL-10 in the intestine following *C. difficile* infection. **(A)** Relative proportion of IL-10-GFP^+^ cells in the large intestine lamina propria (LI Lp) of uninfected and *C. difficile* infected *Il10^GFP^* mice at Day 21 p.i. (Day 0 post-FMT). **(B)** Frequency and **(C)** total number of Foxp3^+^ IL-10-GFP^+^ CD4^+^ T cells in the LI Lp of uninfected and *C. difficile* infected *Il10^GFP^* mice at Day 21 p.i. Gating strategy shown in Supplemental **Fig S2A**. **(D)** Mean fluorescence intensity (MFI) of transcription factors and co-inhibitory receptors in IL-10-GFP^+^ Foxp3^+^ T_reg_ cells and IL-10-GFP^NEG^ Foxp3^+^ T_reg_ cells from *C. difficile* infected mice at Day 21 p.i. Grey shaded histogram represent Fluorescence Minus One (FMO) controls for Rorγt and T-bet. Statistical significance was calculated by two-sided **(A, C)** unpaired or **(D)** paired *t*-test. Data is representative of 3 independent experiments. **(E)** *C. difficile* burden in fecal pellets from *CD4^Cre^Il10^FL/FL^* (n=6) and *CD4^Cre^* (n=7) mice post FMT. Data is combined from 3 independent experiments. Data are presented as mean values ± SEM. *C. difficile* burden statistical significance was calculated by two-part model *t*-test. * p < 0.05, ** p < 0.01, *** p < 0.001. L.o.D.-Limit of Detection.

CX3CR1 macrophages are a minor contributor to the overall pool of IL-10 expressing cells during *C. difficile* infection (**Fig 2A**, **Suppl Fig 2B**). However basal IL-10 production from CX3CR1 macrophages can help limit colonic inflammation in other diseased settings^64,65,64,65^ To confirm that IL-10 from CD4^+^ T cells and not IL-10 derived from myeloid cells was necessary to support FMT treatment, *Cd4*^Cre^ *Il10*^Flox/Flox^ mice along with *Cd4*^Cre^ control mice were generated, infected with *C. difficile,* and administered an FMT. *Cd4*^Cre^ *Il10*^Flox/Flox^ mice exhibited significantly impaired resolution of *C. difficile* following FMT compared to *Cd4*^Cre^ control mice demonstrating that IL-10 from the T cell lineage is needed to support FMT efficacy (**Fig. 2E**). Combined these data support a model where IL-10 predominantly from T_reg_ cells plays a critical role in maintaining immune homeostasis during *C. difficile* infection thereby establishing an environment supportive of FMT treatment.

### T_reg_ cell depletion leads to IFNγ induction

The inability of IL-10 deficient or T_reg_ cell depleted mice^26^ to resolve *C. difficile* infection following FMT suggests an unregulated inflammatory immune response directly inhibits FMT efficacy. Therefore, the downstream cellular and cytokine network regulated by T_reg_ cells that shape FMT-mediated resolution of *C. difficile* were investigated. T_reg_ cell depleted, *C. difficile* infected *Foxp3^DTR^* mice exhibit a broad increase in intestinal inflammatory markers as defined by elevated Lipocalin-2 level in the feces (**Suppl. Fig. 3A**), increased neutrophil infiltration into the large intestinal lamina propria, and upregulation of inflammatory cytokines^26^. *Ex vivo* stimulation of the large intestinal lamina propria cells isolated from T_reg_ cell depleted *C. difficile* infected mice on the day of FMT (day 21 p.i.) demonstrated that both CD4^+^ T cells (**Fig. 3A**) and Innate Lymphoid Cells (ILCs) (**Fig. 3C**) have increased IFN-γ production capacity in both frequency (**Fig. 3A, C**) and total cell numbers (**Fig. 3B, D**) compared to *C. difficile* infected T_reg_ cell sufficient mice, while no induction of IL-17A competent cells was observed (**Fig. 3E**, **F**). As early as 4 days post T_reg_ cell depletion an elevated *Ifng*-*Nos2* signature was observed in the colon, and it correlated with elevated Lipocalin-2 (**Fig. 3G, H**) and increased neutrophils (**Fig. 3I, J**) in the intestine provoking a hypothesis that this immune pathway induces a cascade of inflammatory events leading to FMT failure. *Ex vivo* stimulation of the large intestinal lamina propria cells at this early timepoint following T_reg_ cell depletion confirmed that both CD4^+^ T cells (**Suppl. Fig. 3B, C**) and ILCs (**Suppl. Fig. 3D, E**) rapidly increased their capacity to produce IFN-γ. The IFN-γ signature was unique to mice that were both infected with *C. difficile* and depleted of T_reg_ cells (**Fig. 3G, I**), indicating that the loss of both immunoregulatory networks and the inflammatory response to an ongoing infection were needed to drive strong IFN-γ expression.

**Fig3:**
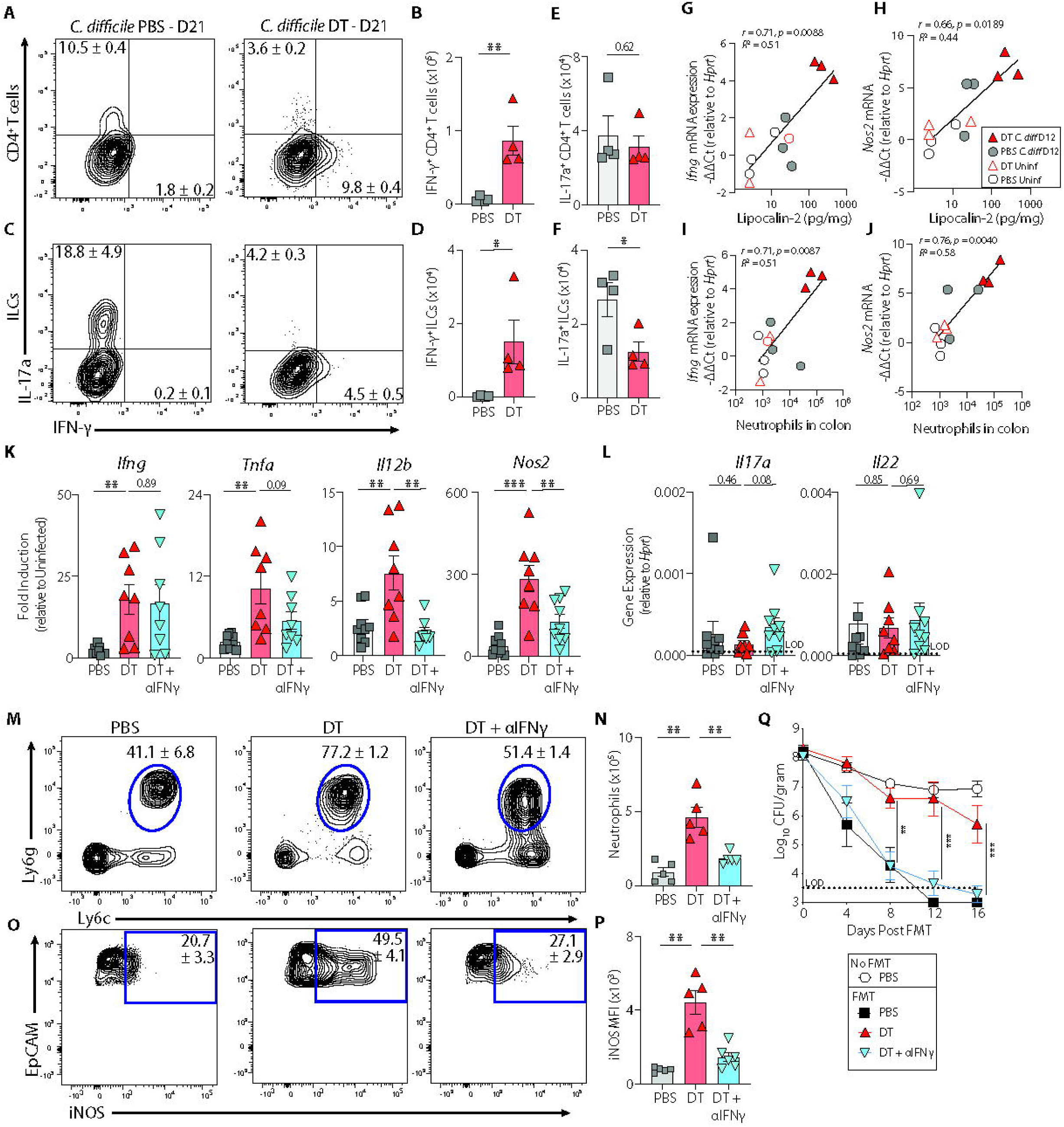
Loss of Treg cells induce IFNγ driven intestinal inflammation but IFNγ blockade restores FMT mediated *C. difficile* resolution (A-F) *Foxp3^DTR^* mice were infected with *C. difficile*, administered (Diphtheria toxin) DT or PBS at Day 8 and 9 p.i. and sacrificed at Day 21 p.i. (Day 0 post-FMT), and single cell suspensions isolated from LI Lp were stimulated with PMA/Ionomycin in the presence of BFA and assessed for cytokine production. **(A)** Frequency and total number of **(B)** IFNγ^+^ CD4^+^ T cells and **(E)** IL-17a^+^ CD4^+^ T cells. FACS plot gated on Live, CD45^+^, CD3/5^+^, CD4^+^ T cells. **(C)** Frequency and total number of **(D)** IFNγ^+^ and **(F)** IL-17a^+^ Innate Lymphoid Cells (ILCs). FACS plot gated on Live, CD45^+^, CD3/5^NEG^, CD19^NEG^, Gr-1^NEG^, CD90^+^ cells. **(G-J)** *C. difficile* infected and uninfected *Foxp3^DTR^* mice were administered DT or PBS at Day 8 and 9 p.i. and sacrificed at Day 12 p.i (n=3 mice/group). Pearson correlation between fecal Lipocalin-2 concentration and **(G)** *Ifnγ* and **(H)** *Nos2* log2 fold mRNA gene expression in colon tissue 4 days post T_reg_ cell depletion (Day 12 p.i.). Pearson correlation between total neutrophils numbers in LI Lp and (**I**) *Ifnγ* and **(J)** *Nos2* log2 fold mRNA gene expression in colon tissue 4 days post T_reg_ cell depletion (Day 12 p.i.). **(K-P).** *Foxp3^DTR^* mice were infected with *C. difficile*, administered DT or PBS at Day 8 and 9 p.i., treated with anti-IFN-γ or αIgG every 3 days starting at Day 12 p.i. and sacrificed at Day 21p.i. (Day 0 post-FMT) to assess immune status. **(K)** Fold induction of *Ifng* associated immune response genes in colon tissue relative to ABX-treated, uninfected *Foxp3^DTR^* mice and normalized to *Hprt* gene. **(L)** mRNA gene expression of *Il17a* and *Il22* in colon tissue relative to *Hprt*. Gene expression of *Il17a* and *Il22* in ABX-treated, uninfected *Foxp3^DTR^* mice were below limit of detection, therefore gene expression is displayed as relative to *Hprt.* Data is combined from 2 independent experiments. **(M)** Frequency and **(N)** total number of neutrophils in LI Lp. FACS plot gated on live, CD45^+^ non-T and non-B, CD11b^+^ cells. **(O)** Frequency and **(P)** mean florescence intensity (MFI) of iNOS expressing large intestinal epithelial cells. FACS plot gated on live, CD45^NEG^, EpCAM^+^ cells. Gating strategy shown in Supplemental **FigS3G**. **(Q)** *C. difficile* burden post FMT, in fecal pellets from *Foxp3^DTR^* mice treated with PBS + αIgG (n = 7), DT + αIgG (n = 7), DT + αIFN*γ* (n = 8) antibodies and PBS treated mice not receiving FMT (n = 7). Data is combined from 2 independent experiments. Experimental design shown in Supplemental **Fig S3F**. Data are presented as mean values ± SEM. Statistical significance was calculated by two-sided unpaired *t*-test. *C. difficile* burden statistical significance was calculated by two-part model *t*-test. * p < 0.05, ** p < 0.01, *** p < 0.001. L.o.D.-Limit of Detection.

To test the hypothesis that IFN-γ driven inflammation contributes to FMT failure to resolve *C. difficile* infection, IFN-γ neutralizing antibodies or isotype IgG control antibodies were administered to T_reg_ cell depleted mice prior to FMT (**Suppl. Fig 3F**). On the day of FMT, *Ifng* expression in the colon remained elevated in *C. difficile* infected, T_reg_ cell depleted mice but IFN-γ blockade had reduced expression of the *Ifng* responsive gene *Nos2*, *Il12b,* and *Tnfa* (**Fig. 3K**). In contrast, colonic *Il17a* and *Il22* expression remained unchanged in T_reg_ cell depleted mice regardless of IFN-γ blockade therapy (**Fig. 3L**). IFN-γ blockade also decreased neutrophil infiltration in the large intestine (**Fig. 3M, N**), and decreased the frequency (**Fig. 3O**) and intensity (**Fig. 3O, P, Suppl. Fig. 3G**) of iNOS expression in EpCAM^+^ intestinal epithelial cells. Following FMT, IFN-γ blockade was sufficient to revert FMT non-responsive mice back to FMT Responders and resolve *C. difficile* from the intestine while T_reg_ cell depleted mice treated with isotype control Rat IgG failed to resolve *C. difficile* following FMT (**Fig. 3Q**). Combined these data support a model where loss of immune suppression via T_reg_ cells leads to elevated IFN-γ signaling and the resulting inflammatory response impairs a host’s FMT responsiveness.

### ROS/RNS derivatives inhibit FMT-mediated resolution of *C. difficile* infection

Reactive oxygen species (ROS) and reactive nitrogen species (RNS) derivatives promote the growth of inflammation tolerant bacteria over inflammation sensitive bacteria in the intestinal lumen^66–74^. We hypothesized that increased iNOS expression and ROS/RNS producing neutrophils in T_reg_ cell depleted, *C. difficile* infected mice promote the growth of inflammation tolerant bacteria and prevents engraftment of beneficial inflammation sensitive microbes from the FMT inoculum. Therefore, the contribution of ROS and RNS derivatives to inhibit FMT efficacy was next assessed. First, *E. coli* WT Nissle and a mutant *E. coli* deficient in the *moaA* gene, which is required for the bacteria to use ROS/RNS derivatives as electron acceptors^71,79,80^, were used as *in vivo* bio-indicators to determine if the intestinal luminal environment of T_reg_ cell depleted, *C. difficile* infected mice was enriched in ROS/RNS derivatives and preferentially supports the growth of inflammation-tolerant bacteria. T_reg_ cell depleted mice exhibited significantly elevated fecal Lipocalin-2 compared to T_reg_ cell sufficient mice at day 15 p.i. indicating an intestinal environment more conducive to microbes that tolerate oxidative stress (**Fig. 4A**). Therefore *C. difficile* infected, T_reg_ cell depleted and T_reg_ cell sufficient mice were inoculated at day 15 post infection with a 1:1 ratio of wild-type *E. coli* and *moaA* mutant *E. coli* and competitive growth index between the two *E. coli* strains was assessed in both hosts at day 21 p.i. (day of FMT) (**Suppl. Fig 4A**). Wild-type *E. coli* had a mean 144-fold competitive growth advantage compared to *moaA* mutant *E. coli* in *C. difficile* infected, T_reg_ cell depleted mice. Further, the absolute number of WT *E. coli* in the cecal content was a mean 355-fold higher in T_reg_ cell depleted mice compared to T_reg_ cell sufficient mice (**Fig. 4B**). This result confirms that intestinal lumen of *C. difficile* infected, T_reg_ cell depleted mice harbor ROS/RNS derivatives that preferentially promote the expansion of inflammation tolerant bacteria. To test whether elevated intestinal ROS/RNS impair FMT mediated resolution of *C. difficile* infection, neutrophils, which are high ROS/RNS producers ^75–78^, were depleted in *C. difficile* infected, T_reg_ cell depleted mice using an anti-Ly6g antibody and subsequently administered an FMT (**Suppl. Fig. 4B, C**). Following FMT, anti-Ly6g antibody treated mice had significantly improved resolution of *C. difficile* compared to T_reg_ cell depleted mice administered an isotype control antibody (**Fig. 4C**). Multiple cell types in the intestine are capable of producing ROS/RNS, including intestinal epithelial cells (**Fig. 3O, P**). Therefore, a complementary pharmacological approach was taken to inhibit ROS/RNS production by using aminoguanidine hydrochloride (AG [NOS2 inhibitor])^71,80^ and N-acetyl-cysteine (NAC [ROS inhibitor])^81,82^. AG+NAC were administered in the drinking water of *C. difficile* infected, T_reg_ cell depleted mice and administered an FMT (**Suppl. Fig 4D**). T_reg_ cell depleted mice that received AG+NAC displayed a significantly faster resolution of *C. difficile* infection compared to the T_reg_ cell depleted untreated mice following FMT (**Fig. 4D**). To determine whether ROS/RNS production contributed to FMT failure in the context of IL-10 ablation (**Fig. 1**), *C. difficile* infected C57BL/6 mice were administered anti-IL-10R antibody with or without AG+NAC treatment prior to FMT (**Suppl. Fig 4E**). Mice that received anti-IL-10R and AG+NAC exhibited a significantly decreased *C. difficile* burden following FMT compared to mice receiving anti-IL-10R blockade alone (**Fig. 4E**) with an increased portion of mice completely resolving the infection (**Suppl. Table 2**). Together, these data suggests that FMT mediated resolution of *C. difficile* is dependent on T_reg_ cells and IL-10 orchestrating an immune regulatory network that restrains an inflammatory cascade culminating in increased ROS/RNS derivatives in the intestinal lumen that selectively inhibit engraftment of beneficial bacterial taxa from the FMT.

**Fig4:**
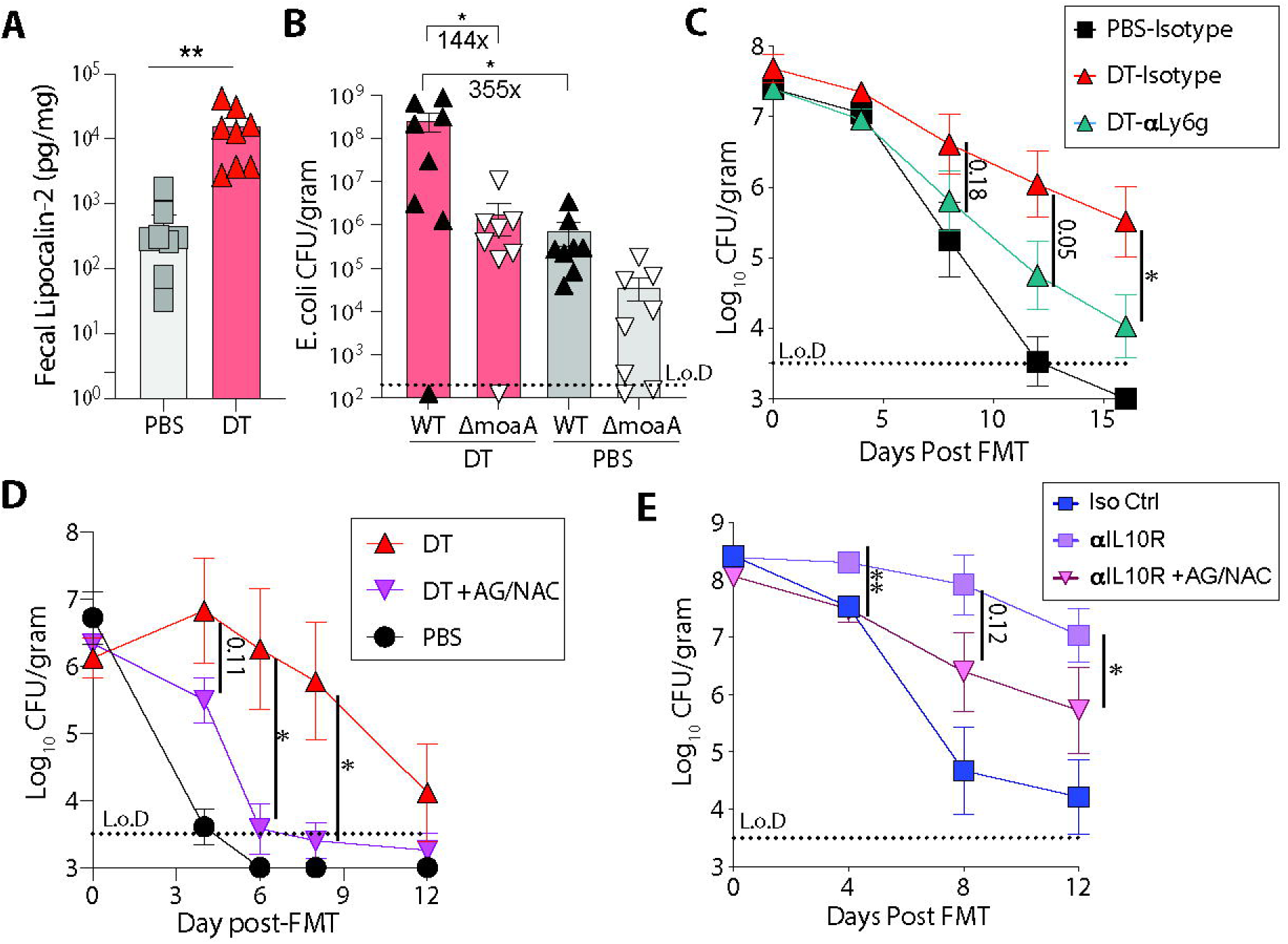
ROS/RNS derivatives inhibit FMT-mediated resolution of *C. difficile* infection. **(A)** Fecal Lipocalin-2 levels in *C. difficile* infected, PBS or DT treated *Foxp3^DTR^* mice on day of *E. coli* administration (Day 15 p.i.). **(B)** Burden of *E. coli* WT Nissle and *E. coli ΔmoaA* mutant bacteria in cecal content from *C. difficile* infected, PBS or DT treated *Foxp3^DTR^*mice at Day 21 p.i. (Day of FMT) n=8 mice/group. Data is combined from two independent experiments. Experimental design shown in Supplemental Fig **S4A**. **(C)** *C. difficile* burden post FMT in fecal pellets from *Foxp3^DTR^* mice treated with PBS + αIgG (n=10), DT + αIgG (n=11) and DT + αLy6g (n=12). Data is combined from two independent experiments. Experimental design shown in Supplemental **Fig S4B**. **(D)** *C. difficile* burden post FMT in fecal pellets from *Foxp3^DTR^* mice treated with PBS (n= 5), DT (n = 4), DT+AG+NAC (n=7). Experimental design shown in Supplemental Fig **S4D**. **(E)** *C. difficile* burden post FMT in fecal pellets from C57BL/6 mice treated with αIgG (n = 10), αIL-10R (n = 10), and αIL-10R+AG+NAC (n = 13). Data is combined from two independent experiments. Experimental design shown in Supplemental **Fig S4E**. Data are presented as mean values ± SEM. Statistical significance was calculated by two-sided unpaired *t*-test for **Fig A,** and both unpaired and paired *t*-test for **Fig B**. *C. difficile* burden statistical significance was calculated by two-part model *t*-test. * p < 0.05, ** p < 0.01, *** p < 0.001. L.o.D.-Limit of Detection.

### *Il10-*mRNA-LNP immunotherapy converts FMT Non-Responders to Responders

Having established that loss of key immunoregulatory factors (T_reg_ cells and IL-10) were sufficient to convert an FMT responsive host into a Non-Responder, we sought to develop an immunotherapy capable of restoring host responsiveness to an FMT. Therefore, we developed an *Il10-*mRNA-LNP platform (**Suppl. Fig. 5A**) to deliver IL-10 protein to FMT Non-Responders and assessed the therapeutic potential of cytokine supplementation immunotherapy to augment FMT efficacy. *In vitro* studies confirmed that the *Il10-*mRNA-LNP induced transcription of biologically active IL-10 protein (**Suppl. Fig. 5B**) capable of enhancing survival of MC/9 cells, an IL-10 responsive cell line^83,84^ (**Suppl. Fig 5C)**. Next, the *in vivo* activity was confirmed by detecting circulating IL-10 protein in the serum of *Il10*^-/-^mice 24 hours following injection with the *Il10-*mRNA-LNP (**Suppl. Fig. 5D**). Finally, intraperitoneal administration of *Il10-*mRNA-LNP into naïve C57BL/6 mice resulted in detectable IL-10 protein in the serum, liver and cecal tissue homogenate for up to 72 hours post administration, peaking at 24 hours, demonstrating *Il10-*mRNA-LNP immunotherapy delivered IL-10 to the large intestinal tissue (**Fig. 5A**). Next, *Il10-*mRNA-LNP was administered every 72 hours to *C. difficile* infected, T_reg_ cell depleted mice starting one day prior to T_reg_ cell depletion and continued throughout the course of FMT administration to ensure sustained IL-10 levels (**Suppl. Fig. 5E**). *Il10-*mRNA-LNP administration resulted in a significant increase in IL-10 levels in the serum at the time of FMT (**Fig. 5B**). *Il10-*mRNA-LNP treated, T_reg_ cell depleted mice exhibited significantly decreased *C. difficile* burden following FMT (**Fig. 5C**) and histologically displayed reduced intestinal epithelial damage (**Fig. 5D**) compared to T_reg_ cell depleted mice treated with an empty vector LNP. Weighted UniFrac (**Fig. 5E**, **F**) and unweighted UniFrac **(Suppl. Fig. 5F, G)** principal coordinate analyses and distance comparisons revealed that there was significantly improved FMT engraftment in *Il10*-mRNA-LNP treated mice compared to empty LNP treated mice. These data provide a proof-of-concept that an immunotherapy that targets a specific immune pathway can be used in combination with a microbiome-based therapeutic to convey treatment success when either therapeutic approach in isolation results in treatment failure.

**Fig5:**
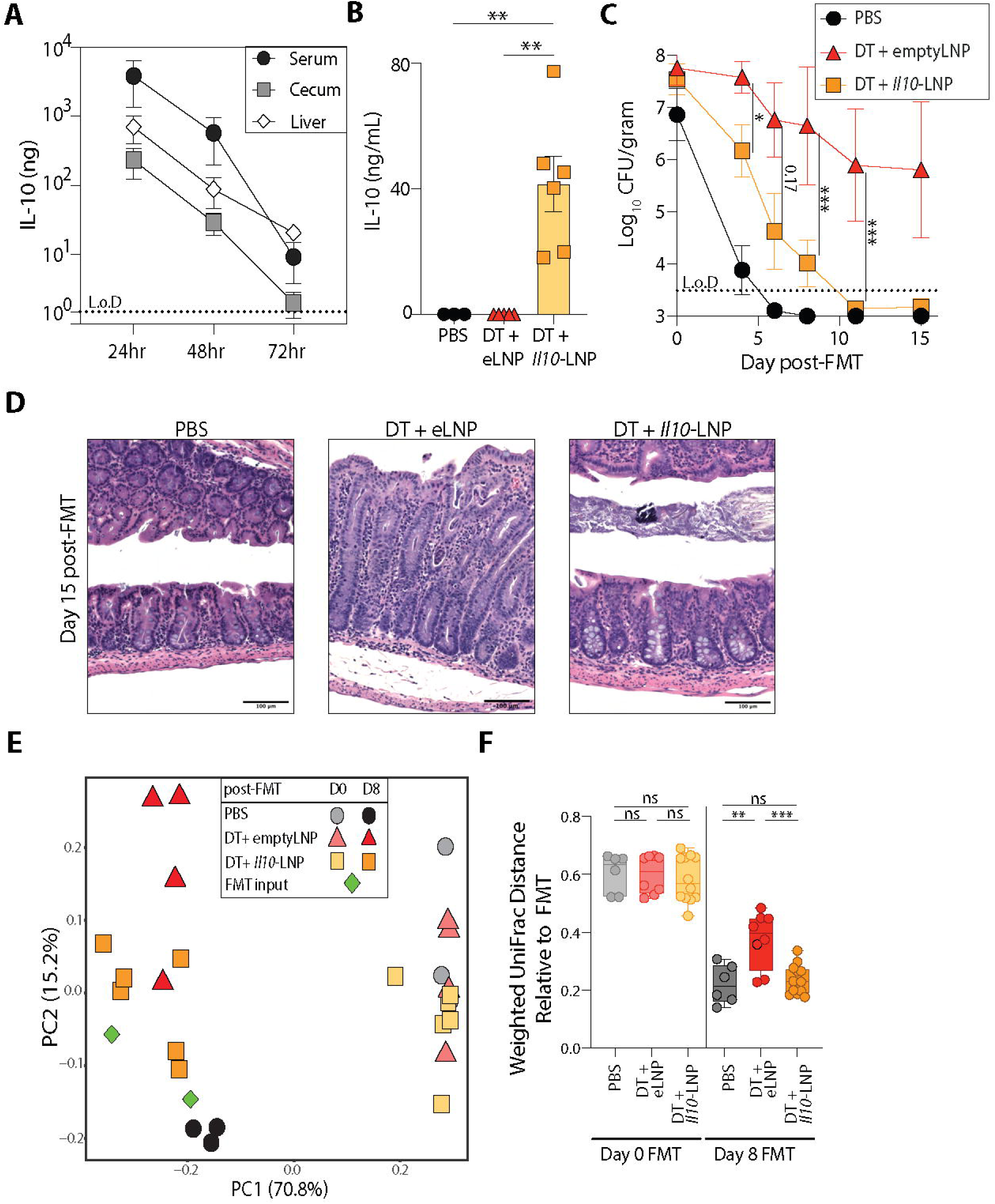
Il10mRNA-LNP immunotherapy converts FMT Non-Responder mice to Responders. **(A)** IL-10 protein level in serum (ng/mL), cecum and liver tissue homogenate (ng/mg) from naïve C57BL/6 mice at different timepoints post intraperitoneal *Il10*mRNA-LNP (10 μg) administration. n=3 mice/timepoint. **(B-F)** *C. difficile* infected *Foxp3^DTR^* mice were administered DT or PBS and treated with empty-LNP (eLNP) or *Il10-*mRNA-LNP (*Il10*-LNP) every 72 hours starting at Day 7 p.i. and administered an FMT at Day 12 p.i.. Experimental design shown in Supplemental **Fig S5E**. PBS (n = 3), DT+ empty-LNP (n = 4) and DT+*Il10-*mRNA-LNP (n = 6). Data representative of two independent experiments. **(B)** IL-10 protein level in serum at Day 12 p.i. (Day of FMT). (**C)** *C. difficile* burden post FMT. **(D)** Representative hematoxylin and eosin stained cecal tissue section at day 15 post-FMT. **(E)** Weighted UniFrac principal coordinate analysis plot of 16S bacterial rRNA ASVs from fecal pellets of PBS, DT+eLNP and DT+*Il10*-LNP groups at Day 0 and 8 post-FMT. **(F)** Weighted UniFrac distance comparing the microbial beta diversity dissimilarity of PBS, DT+eLNP and DT+*Il10*-LNP groups to the FMT inoculum. Data are presented as mean values ± SEM. Statistical significance was calculated by two-sided unpaired *t*-test. *C. difficile* burden statistical significance was calculated by two-part model *t*-test. UniFrac distance statistical significance was calculated by one-way ANOVA test. * p < 0.05, **p <0.01, *** p < 0.001. Boxes represent median and the first and third quartile. Whiskers extend to the highest and lowest data point. L.o.D.-Limit of Detection.

## Discussion

Restoration of beneficial, diverse microbial communities, not only re-establishes colonization resistance against pathogens but also improves gut barrier function and dampens proinflammatory responses, as demonstrated in multiple animal studies^17–19,85,86^. FMT therapy has yielded considerable success in treating patients with uncomplicated, recurrent *C. difficile* infection^4,6,34^. However, in cases where the host’s immune status is overactivated and unchecked, such as in severe, complicated *C. difficile*, or IBD flares, FMT does not have a consistent record of success^35–38^. Current FMT strategies largely adopt a “one-size-fits-all” approach, with limited consideration of the host’s underlying immune landscape^87^. A deeper understanding of host–microbiome–immune interactions prior to FMT could therefore enable a more precision-based framework, guiding patient stratification and optimizing treatment strategies. Such an approach may help identify which patients are most likely to benefit from FMT, who may require adjunctive immunomodulatory therapies, and whether repeated or tailored microbial interventions are necessary to achieve sustained clinical benefit.

In this report, we demonstrate that mice experiencing intestinal inflammation via loss of IL-10 signaling, either due to genetic deletion or antibody blockade, fail to engraft the microbes of a fecal transplant and cannot resolve *C. difficile* infection. Our group previously established a role for the host immune system, specifically CD4^+^ T_reg_ cells, in support of successful FMT mediated resolution of *C. difficile* infection^26^. Here we report that responsiveness can be therapeutically manipulated, and FMT Non-Responders can be converted into Responders by modulating immune pathways. Loss of immune suppression by T_reg_ cells via IL-10 secretion results in increased pro-inflammatory IFN-γ expression in the large intestine that drives neutrophil recruitment and elevated ROS/RNS production. Previous reports have shown that ROS/RNS derivatives promote growth of inflammation tolerant bacteria^66–74^ that negatively influence disease outcomes. Similarly, we observed that the intestinal lumen of FMT non-responsive mice promoted the growth of inflammation tolerant bacteria and impeded the engraftment of microbes from the transplant. Targeting the IFN-γ pathway, neutrophils or production of ROS/RNS all restored FMT’s capacity to resolve *C. difficile* infection. Lastly, we show that an *Il10-*mRNA-LNP immunotherapy was sufficient to enable FMT engraftment and convert Non-Responder mice into Responders.

IL-10 supplementation as a therapeutic approach to treat inflammatory disease has an extensive history of success in animal models^88–91^ that has not translated to the clinic^92,93^. Potential limitations of previous approaches are the inability to deliver recombinant IL-10 protein locally to the desired tissue site, short half-life of the IL-10 bioactivity following injection, and prohibitive cost of administering recombinant protein at the frequency and quantity necessary to deliver a therapeutic effect. Using mRNA-LNP technology can begin to address these issues and make cytokine supplementation therapy viable. Indeed, delivery of *Il12-*mRNA-LNP was found to augment the magnitude and quality of vaccine induced CD8^+^ T cell responses^94^, and *Il22-*mRNA-LNP led to healing of mucosal injury and reduced pro-inflammatory cytokines levels in mice^95^. More specifically to IL-10, an *Il10-*mRNA platform has been used to provide therapeutic benefits in mouse models of IBD, intracerebral hemorrhages, spinal cord injury, and solid tumors^96–99^. In this report we demonstrate for the first time that a cytokine supplementation approach using mRNA-LNP technology is a promising option to support microbiome-based therapeutics. The *Il10-*mRNA-LNP delivered high levels of IL-10 cytokine both systemically and to the intestine for up to 72 hours. *Il10-*mRNA-LNP treatment by itself did not lead to resolution of *C. difficile* infection; however, in conjunction with an FMT, mice exhibited resolution of *C. difficile* driven intestinal inflammation. Although we observe promising therapeutic benefits from this *Il10-*mRNA-LNP formulation, there are several limitations with this current platform that require further optimization to improve the overall efficiency of this treatment. First, multiple 10 μg doses of the *Il10-*mRNA-LNP were administered to maintain elevated IL-10 levels in the intestine over the course of the FMT. Repeated treatments have the potential to result in adverse side effects due to the immunostimulatory nature of the LNP itself, that may counteract the overall immunoregulatory benefits of IL-10. In addition, intraperitoneal delivery of *Il10-*mRNA-LNP induced systemic IL-10 and therefore was not a localized intestinal immunoregulatory therapy. In some therapeutic scenarios broad immunosuppression induced by systemic IL-10 may dampen host defense capabilities and leave the host vulnerable to opportunistic pathogens at other sites of the body. An *Il10-*mRNA-LNP platform designed to deliver IL-10 to the specific cell type or tissue location could provide a more targeted approach that would improve delivery and thereby reducing the amount and frequency of treatments. These caveats notwithstanding, our study demonstrates that mRNA-LNP technology to deliver cytokine therapy is a promising approach to augment microbiome-based therapeutics.

The scientific principles underlying microbiome-based therapeutics as a medical treatment is that the transplanted microbes will convey beneficial properties to the recipient diseased host. Concurrently modulating host factors that condition the recipient patients for successful engraftment of a microbial consortia remains an untapped component that can increase success rates of microbiome-based therapeutics. In this context, the state of microbiome transplantation therapy is analogous to the early years of solid organ transplantation. Similar to advances made in Transplant Immunology over the past sixty years to identify immune activation pathways and develop immunosuppressive drugs to reduce the risk of solid organ transplant rejection, identifying and targeting specific immunological pathways that support microbiome-based transplant engraftment may be necessary for a broader adoption of this treatment strategy. In this report we highlight the IL-10-IFNγ-ROS/RNS axis as a potential target to improve FMT therapy to treat *C. difficile* infection. However, the specific immune pathways to target for each microbiome therapy may be determined by the disease being treated, composition of microbial transplant and the immune status of the microbiome transplant candidate prior to transplantation. This report provides proof-of-concept that a combinatorial immunotherapy-microbiome therapeutic approach can increase the efficacy of microbiome-based therapeutics to treat disorders driven by an underlying microbiome dysbiosis.

## Methods

### Mice

All mice strains were derived on a C57BL/6 background. *Il10^-/-^, Il10^GFP^ (Vert-X), Foxp3^DTR^*, *Cd4*^Cre^, *Il10*^Flox/Flox^ mice were purchased from the Jackson Laboratory. *Cd4*^Cre^ *Il10*^Flox/Flox^ mice were generated by crossing *CD4^Cre^* mice with *Il10^Flox/Flox^* mice. All mice were bred and maintained in sterile autoclaved cages under specific pathogen-free conditions and kept on a grain-based diet (Labdiet 5010) at the University of Pennsylvania. Mice were provided autoclaved water ad libitum from water bottles. Age matched (Eight-to sixteen-week-old) mice of both sexes, with appropriate littermate controls were used in experiments according to institutional guidelines for animal care. To ensure similar starting microbiome composition, mice in experimental groups were cohoused before treatment. All animal procedures were approved by the Institutional Animal Care and Use Committee of the University of Pennsylvania.

### Antibiotic pretreatment, *C. difficile* infection and mouse monitoring

To induce susceptibility to *C. difficile* infection, mice were administered drinking water supplemented with antibiotics: 0.25 g/L metronidazole (Sigma-Aldrich), 0.33 g/L neomycin (Sigma-Aldrich), and 0.33 g/L vancomycin (Novaplus) for four days, and then replaced with normal water for the duration of the experiment. Forty-eight hours following cessation of antibiotic water, mice received 200 μg of clindamycin (Sigma-Aldrich) by intraperitoneal injection (i.p.). Twenty-four hours later, mice received approximately 1000 *C. difficile* spores (CD196 strain) via oral gavage (**Supple.** **Fig 1A**). Mice that exhibited severe disease, defined as surface body temperature below 29.5°C and weight loss greater than 30% were humanely euthanized by CO_2_ displacement.

### Quantification of *C. difficile* burden

Fecal pellets or cecal content were weighed and then resuspended in 1 mL deoxygenated PBS (Corning). Ten-fold serial dilutions were plated overnight at 37°C in an anaerobic chamber (Coylabs) on CCBHISTA agar: 16 g/L agar (BD Biosciences) supplemented with 37 g/L brain heart infusion (BD Biosciences), 5 g/L yeast extract (BD Biosciences), 1 g/L taurocholic acid (MP Biomedicals), 1 g/L L-cysteine (Sigma-Aldrich), 0.25 g/L D-cycloserine (Sigma-Aldrich) and 8 mg/L cefoxitin (Sigma-Aldrich). CFUs were enumerated to calculate the CFU/gram fecal pellet. Prior to infection, fecal pellets of mice were cultured overnight in CCBHISTA liquid broth at 37°C in an anaerobic chamber and then ten-fold serial dilutions were plated on CCBHISTA agar to ensure the absence of endogenous *C. difficile*^107^. Supernatants of fecal or cecal content were obtained after centrifugation for subsequent ELISAs.

### IL-10 & Lipocalin-2 ELISA

Concentration of IL-10 was quantified in cell culture supernatants, serum, liver and cecal tissue homogenates using Mouse IL-10 DuoSet ELISA kit (R&D Systems) as per manufacturer’s instruction. The concentration of Lipocalin-2 in fecal or cecal content supernatant was quantified using DuoSet Lipocalin-2/NGAL ELISA kit (R&D Systems) as per manufacturer’s instruction.

### FMT treatment

The source of the FMT was fecal pellets from naïve, C57BL/6 mice bred in the University of Pennsylvania vivarium and housed under specific pathogen free conditions. Feces were screened for the absence of endogenous *C. difficile* strains^107^. Fecal pellets were collected fresh for each FMT, resuspended at a concentration of 0.2-0.25 grams of feces/mL of deoxygenated PBS (Corning) under anaerobic conditions to preserve obligate anaerobic bacteria. FMT inoculum was flash spun to settle food debris and administered to mice via oral gavage (200 μl) and intrarectal instillation (100 μl) into chronically *C. difficile* infected mice. Twenty-four hours following initial FMT, mice receive a second dose of freshly prepared FMT via oral gavage (200 μl). To normalize FMT across experiments, fecal pellets were collected from the same colony of C57BL/6 mice. Following FMT, mice were separated into individually housed cages.

### T_reg_ cell depletion, antibody blockade and ROS/RNS inhibition

For T_reg_ cell depletion experiments, *Foxp3^DTR^* mice received 500 ng of diphtheria toxin (Sigma-Aldrich) or 200 μl PBS (Corning) as control via intraperitoneal (i.p.) injection on day 8 and 9 post infection (p.i.). For specific cytokine neutralization or immune cell depletion experiments, C57BL/6 mice received via i.p. injection: 1 mg of αIL-10R (clone 1B1.3A; Bio X Cell) once per week for IL-10 blockade, 400 μg of αIFN-γ (clone XMG1.2; Bio X Cell) every 72 hours for IFN-γ blockade, 300 μg of αLygG (clone 1A8; Bio X Cell) every 48 hours for neutrophil depletion respectively. Rat IgG1 (Bio X Cell) was used as isotype antibody control treatment. For ROS/RNS inhibition, C57BL/6 or *Foxp3^DTR^* mice received aminoguanidine hydrochloride (Sigma Aldrich) in fresh drinking water (1 mg/ml) and 2 mg via oral gavage, alongside 20 mg N-acetyl-cysteine (Sigma Aldrich) via oral gavage as indicated in the experimental design.

### Escherichia coli in vivo competition index

*E. coli* Nissle wildtype (pWSK29 Carb^r^) and *E. coli* Nissle Δ*moaA (*pWSK129, Kan^r^) strains were grown overnight (∼16 h) at 37°C in 10 ml Luria Broth (25 gm/L) (BD Biosciences) supplemented with 100 μg/ml Carbenicillin (Sigma) or Kanamycin (Thermoscientfic) respectively. The construction of these bacterial strains(Fu Wang and Kushner, 1991; Winter *et al.*, 2013a)ner, 1991; Winter et al., 2013a). Cells from overnight cultures were harvested by centrifugation (4000 rpm for 5 mins) and resuspended in PBS. The strains were mixed at 1:1 ratio based on their OD_600nm_ measurements. *C. difficile* infected, DT or PBS treated *Foxp3^DTR^* mice received approximately 10^8^ CFU of each bacterium via oral gavage at day 15 p.i. (day 7 post DT administration). At day 21 p.i. cecal content were collected, weighed and resuspended in 1 mL deoxygenated PBS. Ten-fold serial dilutions were plated on LB-Agar (16 g/L) plates supplemented with 100 μg/ml Carbenicillin or Kanamycin to enumerate the respective bacterial strains. CFU/gram of fecal pellet for each strain was calculated to measure the competitive growth index of the two *E. coli* strains.

### *Il10-mRNA*-LNP platform design and *in vitro* and *in vivo* validation

Sequence for mouse IL-10 was codon optimized and cloned into our transcription plasmid template containing a T7 promoter, 5’ and 3’ UTRs and a poly A (100 nucleotide) tail. mRNA was synthesized by *in vitro* transcription incorporating m1ψ-5’-triphosphate (pseudouridine) using Megascript transcription kit (Thermo, AMB 1334) and co-transcriptionally capped with CleanCap^TM^ trinucleotide system (Trilink Biotechnologies). mRNA was precipitated, purified using modified cellulose-based method, resuspended in nuclease- and pyrogen-free water and stored at-20°C until formulation. Before formulation, the mRNA integrity was verified using the Agilent Bioanalyzer 2100 system. Dot blot was used to confirm the removal of double-stranded (ds) RNA. mRNA was encapsulated into ALC-307 (proprietary ionizable lipid of Acuitas Therapeutics) using a self-assembly process where an ethanolic mixture of the ionizable lipid, cholesterol, DSPC and PEG was mixed rapidly with an aqueous solution of the mRNA followed by dialysis and concentration to 1 mg/ml. The hydrodynamic size and polydispersity index (PDI) were 70-80 nm and 0.05-0.13 respectively as determined using dynamic light scattering. Ribogreen RNA assay was used to measure the encapsulation efficiency (97-98%) and concentration of the LNPs (ALC-307). The LNPs were stored at -80°C until use.

*In vitro* validation of *Il10*-mRNA-LNP was done by transfecting HEK293T cells grown at 37°C in 5% CO_2_ in complete TC media (Dulbecco’s Modified Eagle Medium (DMEM, Corning) supplemented with sodium pyruvate (Sigma Aldrich) + 10% FBS (GeminiBio) + 200 mM L-glutamine (Corning) + penicillin-streptomycin (Gibco)) with increasing doses of *Il10*-mRNA-LNP or empty-LNP control. Twenty-four hours later media from transfected wells were collected and IL-10 ELISA conducted. To assess biological activity of transcribed IL-10, MC/9 cell line was grown at 37°C in 5% CO_2_ in complete TC media. MC/9 cells were seeded on 96 well flat bottom plate and transfected with increasing doses of *Il10*-mRNA-LNP or empty-LNP. Twenty-four hours later MC/9 cells per well were enumerated by trypan blue staining and counted on hemocytometer. For *in vivo mRNA*-LNP experiment mice received 10 μg of *Il10*-mRNA-LNP or empty-LNP via i.p. or intramuscular injection as indicated in the experimental design.

### Bacterial DNA extraction, 16S rRNA sequencing and analysis

DNA was extracted from fecal pellets or cecal content with Qiacube (Qiagen) using the DNeasy Powersoil Pro Kit (Qiagen) as per the manufacturer’s instruction. Extracted DNA samples were sent to PennCHOP Microbiome Core, Philadelphia, USA, for 16S rRNA sequencing. Amplicons of the V1-V2 16S rRNA region were amplified and sequenced using an Illumina MiSeq platform as described previously^109^. Sequenced data were imported into QIIME2 (v.2024.10)^110^ and denoised using the DADA2 plugin^111^. Resulting data were taxonomically classified in QIIME2 using the q2-greengenes2^112^ function and comparing against the full-length backbone^113^. Phylogenetic trees were generated using mafft and q2-phylogeny plugin^112^. Data were then imported into R 4.4.2^114^ for further analyses with phyloseq (v.1.30.0)^115^ and visualization with ggplot2 (v.3.3.0)^116^. Unweighted and weighted UniFrac^117^ dissimilarity was calculated to generate principal coordinate analysis (PCoA) plots.

### Tissue processing, RNA isolation, cDNA preparation and RT-PCR

Cecal tissues were fixed with 4% paraformaldehyde, embedded in paraffin and 5 µm sections were cut and stained with hematoxylin and eosin. Tissue sections were then imaged at 40X resolution using a ThermoFisher EVOS M7000 widefield microscope at the Cell and Developmental Biology Microscopy Core. 1-2 cm of the proximal colon was harvested and stored in RNAlater (Invitrogen) at -80°C until RNA extraction was performed. RNA was isolated from proximal colon tissue after mechanical disruption in TissueLyser II (Qiagen) and then in Qiacube (Qiagen) using RNeasy mini kit (Qiagen) according to the manufacturer’s instructions. cDNA was generated using QuantiNova Reverse Transcriptase Kit (Qiagen) as per manufacturer’s instruction. Quantitative RT-PCR was performed on cDNA using the Quantinova SYBR Green PCR Kit (Qiagen) with Quantitect primer assays (Qiagen). Reactions were run on a Quantstudio 6 Flex (Applied Biosystems). Genes of interest were displayed in arbitrary units relative to the expression of *Hprt* in antibiotics (ABX) treated uninfected mice.

### Isolation of cells from lamina propria, *ex vivo* cell stimulation and flow cytometry

Large intestine (cecum and colon) was harvested, opened longitudinally, and contents removed with PBS. Single cell suspensions were obtained by incubating intestinal tissue in epithelial strip buffer (PBS, 5 mM EDTA (Invitrogen), 1 mM dithiothreitol (Fisher Scientific), 4% FBS (GeminiBio), and 10 μg/mL penicillin-streptomycin (Gibco)) while shaking (180 rpm) at 37°C. The supernatant was retained to obtain colonic epithelial cells. The remaining tissue was incubated again in epithelial strip buffer for 20 min while shaking (180 rpm) at 37°C to remove intraepithelial lymphocytes. To obtain immune cells from the lamina propria, the tissue was next digested with 1.25 mg/mL collagenase IV (Worthington Biochemical) and 20 μg/ml DNAse I (Sigma-Aldrich) in complete medium (DMEM (Corning)) supplemented with 10% FBS (GeminiBio), 200 μg/mL L-glutamine (Corning), 1 mM sodium pyruvate (Sigma-Aldrich), 20 mM HEPES (Cytiva), 55 μM 2-mercaptoethanol (Gibco), 50 μg/mL gentamicin sulfate (GeminiBio), and 100 U/mL penicillin (Gibco), 100 μg/ml streptomycin (Gibco)) for 30 min while shaking (180 rpm) at 37°C. Suspensions were passed through a 100 μm strainer, followed by a 40 μm strainer to obtain single cell suspensions. Cells were pelleted at 450 x g for 5 min and then resuspended in 1 mL complete medium. Cells were quantified on a Cellometer Auto 2000 (Nexelcom). For *ex vivo* intracellular cytokine detection, cells were cultured in flat bottom non-TC treated 96-well plates for 4-5 hrs at 37°C in complete medium either in the presence of Brefeldin A (BD GolgiPlug, BD Biosciences), or a mix of Belfredin A, Phorbol 12-myristate 13 acetate (50 ng/mL) (Sigma) and Ionomycin (500 ng/mL) (Sigma). Following *ex vivo* stimulation, or directly following isolation of single cell suspension, cells were transferred to 96-well round bottom plates for cell staining for flow cytometry. Antibodies used for staining cells, intracellular cytokines and transcription factors are listed in **Suppl. Table 3**. Briefly cells were washed with PBS and then stained with LIVE/DEAD Fixable Aqua Dead Cell Stain (Invitrogen) for 20 min at 1:600 dilution in PBS and washed with FACS buffer (PBS, 1% BSA, 50 ug/mL Sodium Azide (Fischer Scientfici)) at 450 x g for 4 min at 4°C, before Fc blocking with 0.1 mg/mL anti-mouse CD16/32 (clone 2.4G2, BD Biosciences) and 0.2 mg/mL rat IgG(Sigma-Aldrich) antibody for 5 min at 4°C. Next, antibodies against surface antigens diluted in FACS buffer were incubated with cells for 30 min at 4°C. Following surface staining, cells were washed twice with FACS buffer at 450 x g for 4 min at 4°C. Thereafter cells were either fixed with 2% paraformaldehyde solution for 15 min at 4°C for surface staining only or proceeded with intracellular cytokine staining or intranuclear staining. For intracellular cytokine staining, cells were treated with eBiosciences intracellular cytokine fixation buffer (Invitrogen) for 30 min at 4°C and washed twice with perm-wash buffer (eBiosciences 1x permeabilization buffer (Invitrogen)) at 450 x g for 4 min at 4°C. Intracellular antibodies diluted in perm-wash buffer were incubated with cells for 30 min at 4°C. Thereafter cells were again washed with perm-wash buffer and then with FACS buffer at 450 x g for 4 min at 4°C before fixation with 2% paraformaldehyde solution for 15 min at 4°C. For intranuclear staining, cells were fixed and permeabilized for 60 min at 4°C using a Foxp3/Transcription factor staining set (eBiosciences) followed by intranuclear antibody staining. Cells were washed twice with perm-wash buffer and once with FACS buffer at 450 x g for 4 min at 4°C and resuspended in FACS buffer. Cells were analyzed on BD LSR-II or Symphony A3 flow cytometer (BD Biosciences) and data was generated in the Penn Cytomics and Cell Sorting Shared Resource Laboratory at the University of Pennsylvania (RRID: SCR_022376). Flow cytometer data was analyzed using FlowJo version 10.10.

### Statistical analysis

Data was collected in Microsoft Excel (version v16.0) and transformed in R (version 4.4.2)^114^ or directly transferred to Graphpad Prism (version 10.0.0) for data visualization and statistical analysis. Results are represented as mean ± standard error of mean (SEM). Statistical significance was determined by unpaired, paired two-sided *t*-test and one-way ANOVA test (* p < 0.05; ** p < 0.01; *** p < 0.001). For *C. difficile* burden where a portion of samples had burden below the limit of detection, values were converted into log_10_ values and then statistical analysis was performed using two-part model *t*-test ^118,119^. For 16S sequencing data, in order to test the null hypothesis of no differences in the study group centroids, a permutation based multivariate ANOVA test as implemented by the function adonis() in the vegan package 2.7-2^120^ was used in R 4.4.2^114^.

FigS1: **Fecal Microbiota Transplantation (FMT) mediated *C. difficile* resolution depends on intact IL-10 signalling prior to FMT**

**(A)** Schematic of antibiotic (ABX) regimen, *C. difficile* infection and Fecal Microbiota Transplantation (FMT). **(B)** *C. difficile* burden in fecal pellets from *Il10^HET^* (n=5) and *Il10^-/-^*(n=4) mice at Day 12 and 21 post infection (p.i.). **(C)** Unweighted UniFrac principal coordinate analysis plot of 16S bacterial rRNA ASVs from fecal pellets of *Il10^HET^*(n =5) and *Il10^-/-^*(n=4) mice at Day 0 and 8 post-FMT. **(D)** Unweighted UniFrac distance comparing the microbial beta diversity dissimilarity of *Il10^HET^* and *Il10^-/-^*groups to the FMT inoculum. **(E, F)** *Il10^HET^* and *Il10^-/-^*mice were treated with the same antibiotic regimen used to predispose mice to *C. difficile* infection but were left uninfected. At 22 days post cessation of antibiotics mice were administered an FMT**. (E)** Weighted UniFrac principal coordinate analysis plot of 16S bacterial rRNA ASVs from fecal pellets of ABX treated uninfected *Il10^HET^* (n =5) and *Il10^-/-^*(n=3) mice at Day 0, 4 and 12 post-FMT. (**F)** Weighted UniFrac distance comparing the microbial beta diversity dissimilarity of *Il10^HET^* and *Il10^-/-^*groups to the FMT inoculum. **(G)** Unweighted UniFrac principal coordinate analysis plot of 16S bacterial rRNA ASVs from fecal pellets of αIgG (n = 3) and αIL-10R (n = 4) groups at Day 0 and 8 post-FMT. (**H)** Unweighted UniFrac distance comparing the microbial beta diversity dissimilarity of αIgG and αIL-10R groups to the FMT inoculum. **(I)** Schematic of ABX regimen, *C. difficile* infection, administration of αIgG and αIL-10R antibodies and FMT. **(J)** *C. difficile* burden in fecal pellets from C57BL/6 mice treated with αIgG (n = 7) or αIL10R antibodies starting either at Day 7 p.i. (n = 8) or Day 21 p.i. (n = 8) mice post FMT. Data is combined from two independent experiments. Data are presented as mean values ± standard error of mean (SEM). *C.difficile* burden statistical significance was calculated by two-part model *t*-test. UniFrac distance statistical significance was calculated by one-way ANOVA test. * p < 0.05, ** p < 0.01, *** p < 0.001. Boxes represent median and the first and third quartile. Whiskers extend to the highest and lowest data point.

FigS2: **IL-10 GFP^+^ Macrophages, B cells and Foxp3^NEG^ CD4^+^ T cells do not expand while IL-10 GFP^+^ T_reg_ cells exhibit a highly immunosuppressive phenotype following *C. difficile* infection**

**(A)** Flow cytometry gating strategy used to identify the immune cell subsets of IL-10 GFP^+^ cells in the large intestine lamina propria (LI Lp). Gating strategy related to **Fig 2A** and Supplementary **Fig S2B**. **(B)** IL10GFP^+^ macrophages, B cells and CD4^+^ T cells of LI Lp of uninfected and *C. difficile* infected *IL10^GFP^*reporter mice. **(C)** Representative FACS plot and mean fluorescence intensity (MFI) of immune markers in IL-10 GFP^+^ Foxp3^+^ T_reg_ cells in uninfected and *C. difficile* infected *IL10^GFP^* reporter mice at Day 21 p.i. Data is representative of 3 independent experiments. Data are presented as mean values ± SEM. Statistical significance was calculated by two-sided unpaired *t*-test. * p < 0.05, **p <0.01, *** p < 0.001.

FigS3: **T_reg_ cell depletion leads to intestinal inflammation with elevated fecal Lipocalin-2 concentration and IFNγ induction**

**(A)** Fecal Lipocalin-2 levels in *C. difficile* infected PBS (n = 14) or DT (n = 16) treated *Foxp3^DTR^* mice at Day 21 p.i. (Day of FMT). Data is combined from three independent experiments. (**B-E)** *Foxp3^DTR^* mice were infected with *C. difficile*, administered DT or PBS at Day 8 and 9 p.i. and sacrificed at Day 12 p.i. (n=3 mice/group). Single cell suspensions isolated from LI Lp were stimulated with PMA/Ionomycin in the presence of BFA and assessed for cytokine production. (**B)** Frequency and (**C**) total number of IFNγ^+^ CD4^+^ T cells. FACS plot gated on Live, CD45^+^, CD3/5^+^, CD4^+^ T cells. (**D)** Frequency and (**E**) total number of IFNγ^+^ ILCs. FACS plot gated on Live, CD45^+^, CD3/5^NEG^, CD19^NEG^, Gr-1^NEG^, CD90^+^ cells. (**F)** Experimental schematic of antibiotic (ABX) regimen, *C. difficile* infection, DT or PBS administration, αIgG or αIFN*γ* antibody administration every 72 hr starting at Day 12 p.i. and FMT at Day 21 p.i. **G.** Flow cytometry gating strategy used to identify EpCAM^+^ epithelial cells and iNOS^+^ large intestinal cells. Gating strategy related to Fig 3O-P. Data are presented as mean values ± SEM. Statistical significance was calculated by two-sided unpaired *t*-test. * p < 0.05, **p <0.01, *** p < 0.001.

FigS4: **Experimental designs for *in vivo E. coli* competition assay and blocking the sources of ROS/RNS**

**(A)** Experimental schematic of *in vivo* competitive growth assay after inoculation with 1:1 ratio of *E. coli* WT Nissle and *E. coli ΔmoaA* mutant. Related to Fig 4A-B**. (B)** Experimental schematic of neutrophil depletion in DT treated *Foxp3^DTR^* mice. αIgG or αLy6g antibody administration every 48 hr starting at Day 12 p.i. Related to Fig. 4C**. (C)** FACS plot showing depletion of neutrophils in LI Lp at Day 21 p.i. (Day of FMT) after αLy6g antibody administration in DT treated *Foxp3^DTR^* mice. FACS plots gated on live, CD45^+^ non-T and non-B, CD11b^+^ cells. (**D)** Experimental schematic of aminoguanidine (AG) and N-acetyl-cysteine (NAC) administration in DT treated *Foxp3^DTR^*mice. Related to Fig 4D. (**E)** Experimental schematic of aminoguanidine (AG) and N-acetyl-cysteine (NAC) administration in C57BL/6 mice treated with αIgG or αIL-10R antibody every 7 days starting at Day 7 p.i. Related to Fig 4E.

FigS5: **Restoration of IL-10 signaling by administration of *Il10-*mRNA-LNP helps in FMT engraftment**

**(A)** Vector map of *Il10-m*RNA-LNP. (**B)** IL-10 protein in supernatants of HEK293T cell line transfected with titrating doses of empty-LNP or *Il10-m*RNA-LNP for 24 hours. n=3 per treatment group. (**C)** Enumeration of MC/9 cells numbers transfected with titrating doses of empty-LNP or *Il10-m*RNA-LNP. n=3 per treatment group (**D)** IL-10 protein level in serum of *Il10*^-/-^mice 24 and 72 hours following administration of *Il10-m*RNA-LNP intraperitoneally (i.p.), intramuscularly (i.m.) or sham mRNA control (n=1 mouse/group). (**E)** Experimental schematic of *Il10-*mRNA-LNP or empty-LNP administration in *C. difficile* infected DT treated *Foxp3^DTR^* mice prior to FMT. (**F)** Unweighted UniFrac principal coordinate analysis plot of 16S bacterial rRNA ASVs from fecal pellets of PBS (n = 3), DT+empty-LNP (n = 4) and *Il10-*mRNA-LNP (n = 6) groups at Day 0 and 8 post-FMT. (**G)** Unweighted Unifrac distance comparing the microbial beta diversity dissimilarity of PBS, DT+empty-LNP and DT+*Il10-*mRNA-LNP groups to the FMT inoculum. Data are presented as mean values ± SEM. Statistical significance was calculated by two-sided unpaired *t*-test. UniFrac distance statistical significance was calculated by one-way ANOVA test. * p < 0.05, ** p < 0.01, *** p < 0.001. Boxes represent median and the first and third quartile. Whiskers extend to the highest and lowest data point. L.o.D.-Limit of Detection.

## Resource availability

### Lead contact

Requests for further information and resources should be directed to and will be fulfilled by the lead contact Michael C. Abt.

### Materials availability

All materials generated in this study will be made available upon reasonable request.

### Data and code availability

- The 16S rRNA gene sequencing data have been deposited at NCBI SRA database as BioProject: PRJNA1416324 and are publicly available as of the date of publication. The source data used to generate figures and full histology images of respective tissue sections will be deposited at FigShare repository and made publicly available as of the date of publication.
- All original code will be deposited at FigShare repository and made publicly available as of the date of publication.
- Any additional information required to reanalyze the data reported in this paper is available from the lead contact upon request.

### Author contributions

S.M, M.Z.A, and M.C.A. contributed to experimental design, performed the experiments and analyzed experimental data. S.M. and M.C.A generated figures and wrote the manuscript. All authors have read, discussed, and edited the manuscript and approved the submitted version. E.N.H, N.V.M, J.R.M, M.T.H., K.S.M, D.D.B assisted in data acquisition. J.E.D. performed 16S rRNA high throughput sequencing analysis. M.G.A provided technical expertise and designed Il10 mRNA plasmid construct.

## Acknowledgements

We thank Dr. Daniel P. Beiting for generously providing us the *E. coli* Nissle wildtype and *E. coli ΔmoaA* mutant strains. We thank Dr. Kevin Amses for his advice on statistical analyses. Flow cytometry data was generated at the Penn Cytomics and Cell Sorting Shared Resource Laboratory of the University of Pennsylvania (RRID: SCR_022376). Tissue sectioning and staining was performed at the Center for Molecular Studies in Digestive and Liver Diseases (P30DK050306) and Molecular Pathology and Imaging Core of the University of Pennsylvania (RRID: SCR_022420). Tissue sections were imaged at the Cell and Developmental Biology Microscopy Core of the University of Pennsylvania (RRID: SCR_022373). *Il10* mRNA-LNP was developed at the Penn Institute for RNA Innovation of the University of Pennsylvania. We thank Dr. Garima Dwivedi from the Penn Institute for RNA mRNA Core for formulation of the *Il10* mRNA-LNP. We would also like to thank all Abt lab members for their helpful suggestions and feedback on the manuscript.

## Abbreviations used

ABX: antibiotic
FMT: Fecal Microbiome Transplant
IBD: Inflammatory Bowel Disease
IL-10: Interleukin 10
ROS: Reactive oxygen species
RNS: Reactive oxygen species
LNP: lipid nanoparticle
T_reg_: Regulatory T cell

## Reference

1. Alam, M.Z., Maslanka, J.R., and Abt, M.C. (2023). Immunological consequences of microbiome-based therapeutics. Front. Immunol. 13, 1–17. 10.3389/fimmu.2022.1046472.

2. Sorbara, M.T., and Pamer, E.G. (2022). Microbiome-based therapeutics. Preprint at Nature Research, 10.1038/s41579-021-00667-9 https://doi.org/10.1038/s41579-021-00667-9.

3. Jain, N., Umar, T.P., Fahner, A.F., and Gibietis, V. (2023). Advancing therapeutics for recurrent clostridioides difficile infections: an overview of vowst’s FDA approval and implications. Preprint at Taylor and Francis Ltd., 10.1080/19490976.2023.2232137 https://doi.org/10.1080/19490976.2023.2232137.

4. van Nood, E., Vrieze, A., Nieuwdorp, M., Fuentes, S., Zoetendal, E.G., de Vos, W.M., Visser, C.E., Kuijper, E.J., Bartelsman, J.F.W.M., Tijssen, J.G.P., et al. (2013). Duodenal Infusion of Donor Feces for Recurrent Clostridium difficile . New England Journal of Medicine 368, 407–415. 10.1056/nejmoa1205037.

5. Rode, A.A., Chehri, M., Krogsgaard, L.R., Heno, K.K., Svendsen, A.T., Ribberholt, I., Helms, M., Engberg, J., Schønning, K., Tvede, M., et al. (2021). Randomised clinical trial: a 12-strain bacterial mixture versus faecal microbiota transplantation versus vancomycin for recurrent Clostridioides difficile infections. Aliment. Pharmacol. Ther. 53, 999–1009. 10.1111/apt.16309.

6. Kelly, C.R., Yen, E.F., Grinspan, A.M., Kahn, S.A., Atreja, A., Lewis, J.D., Moore, T.A., Rubin, D.T., Kim, A.M., Serra, S., et al. (2021). Fecal Microbiota Transplantation Is Highly Effective in Real-World Practice: Initial Results From the FMT National Registry. Gastroenterology 160, 183–192.e3. 10.1053/j.gastro.2020.09.038.

7. Kelly, C.R., Khoruts, A., Staley, C., Sadowsky, M.J., Abd, M., Alani, M., Bakow, B., Curran, P., McKenney, J., Tisch, A., et al. (2016). Effect of fecal microbiota transplantation on recurrence in multiply recurrent clostridium difficile infection a randomized trial. Ann. Intern. Med. 165, 609–616. 10.7326/M16-0271.

8. Doosetty, S., Umeh, C., Eastwood, W., Samreen, I., Penchala, A., Kaur, H., Chilinga, C., Kaur, G., Mohta, T., Nakka, S., et al. (2024). Efficacy of Fecal Microbiota (REBYOTA) in Recurrent Clostridium difficile Infections: A Systematic Review and Meta-Analysis. Cureus. 10.7759/cureus.58862.

9. Feuerstadt, P., Crawford, C. V., Tan, X., Pokhilko, V., Bancke, L., Ng, S., Guthmueller, B., Bidell, M.R., Tillotson, G., Johnson, S., et al. (2023). Fecal Microbiota, Live-jslm for the Prevention of Recurrent Clostridioides difficile Infection Subgroup Analysis of PUNCH CD2 and PUNCH CD3. J. Clin. Gastroenterol. 58, 818–824. 10.1097/MCG.0000000000001947.

10. Sims, M.D., Khanna, S., Feuerstadt, P., Louie, T.J., Kelly, C.R., Huang, E.S., Hohmann, E.L., Wang, E.E.L., Oneto, C., Cohen, S.H., et al. (2023). Safety and Tolerability of SER-109 as an Investigational Microbiome Therapeutic in Adults With Recurrent Clostridioides difficile Infection A Phase 3, Open-Label, Single-Arm Trial. JAMA Netw. Open 6. 10.1001/jamanetworkopen.2022.55758.

11. Feuerstadt, P., Louie, T.J., Lashner, B., Wang, E.E.L., Diao, L., Bryant, J.A., Sims, M., Kraft, C.S., Cohen, S.H., Berenson, C.S., et al. (2022). SER-109, an Oral Microbiome Therapy for Recurrent Clostridioides difficile Infection . New England Journal of Medicine 386, 220–229. 10.1056/nejmoa2106516.

12. Mimee, M., Citorik, R.J., and Lu, T.K. (2016). Microbiome therapeutics — Advances and challenges. Preprint at Elsevier B.V., 10.1016/j.addr.2016.04.032 https://doi.org/10.1016/j.addr.2016.04.032.

13. Yilmaz, O., Okullu, S.O., Catakci, M., Elmas, M.A., Pinheiro, Y., Arbak, S., Demir, E., Schaefer, K.H., and Kolgazi, M. (2024). Akkermansia muciniphila improves chronic colitis-induced enteric neuroinflammation in mice. Neurogastroenterology and Motility 36. 10.1111/nmo.14745.

14. Buffie, C.G., Bucci, V., Stein, R.R., McKenney, P.T., Ling, L., Gobourne, A., No, D., Liu, H., Kinnebrew, M., Viale, A., et al. (2015). Precision microbiome reconstitution restores bile acid mediated resistance to Clostridium difficile. Nature 517, 205–208. 10.1038/nature13828.

15. Lawley, T.D., Clare, S., Walker, A.W., Stares, M.D., Connor, T.R., Raisen, C., Goulding, D., Rad, R., Schreiber, F., Brandt, C., et al. (2012). Targeted Restoration of the Intestinal Microbiota with a Simple, Defined Bacteriotherapy Resolves Relapsing Clostridium difficile Disease in Mice. PLoS Pathog. 8. 10.1371/journal.ppat.1002995.

16. Tian, S., Kim, M.S., Zhao, J., Heber, K., Hao, F., Koslicki, D., Tian, S., Singh, V., Patterson, A.D., and Bisanz, J.E. (2025). A designed synthetic microbiota provides insight to community function in Clostridioides difficile resistance. Cell Host Microbe. 10.1016/j.chom.2025.02.007.

17. Narushima, S., Sugiura, Y., Oshima, K., Atarashi, K., Hattori, M., Suematsu, M., and Honda, K. (2014). Characterization of the 17 strains of regulatory T cell-inducing human-derived Clostridia. Gut Microbes 5, 333–339. 10.4161/gmic.28572.

18. Atarashi, K., Tanoue, T., Oshima, K., Suda, W., Nagano, Y., Nishikawa, H., Fukuda, S., Saito, T., Narushima, S., Hase, K., et al. (2013). Treg induction by a rationally selected mixture of Clostridia strains from the human microbiota. Nature 500, 232–236. 10.1038/nature12331.

19. Round, J.L., and Mazmanian, S.K. (2010). Inducible Foxp3+ regulatory T-cell development by a commensal bacterium of the intestinal microbiota. Proc. Natl. Acad. Sci. U. S. A. 107, 12204–12209. 10.1073/pnas.0909122107.

20. Louie, T., Golan, Y., Khanna, S., Bobilev, D., Erpelding, N., Fratazzi, C., Carini, M., Menon, R., Ruisi, M., Norman, J.M., et al. (2023). VE303, a Defined Bacterial Consortium, for Prevention of Recurrent Clostridioides difficile Infection: A Randomized Clinical Trial. JAMA 329, 1356–1366. 10.1001/jama.2023.4314.

21. Dsouza, M., Menon, R., Crossette, E., Bhattarai, S.K., Schneider, J., Kim, Y.G., Reddy, S., Caballero, S., Felix, C., Cornacchione, L., et al. (2022). Colonization of the live biotherapeutic product VE303 and modulation of the microbiota and metabolites in healthy volunteers. Cell Host Microbe 30, 583–598.e8. 10.1016/j.chom.2022.03.016.

22. Feehley, T., Plunkett, C.H., Bao, R., Choi Hong, S.M., Culleen, E., Belda-Ferre, P., Campbell, E., Aitoro, R., Nocerino, R., Paparo, L., et al. (2019). Healthy infants harbor intestinal bacteria that protect against food allergy. Nat. Med. 25, 448–453. 10.1038/s41591-018-0324-z.

23. Routy, B., Le Chatelier, E., Derosa, L., Duong, C.P.M., Alou, M.T., Daillère, R., Fluckiger, A., Messaoudene, M., Rauber, C., Roberti, M.P., et al. (2018). Gut microbiome influences efficacy of PD-1-based immunotherapy against epithelial tumors. Science (1979). 359, 91–97.

24. Kim, Y.-G., Sakamoto, K., Seo, S.-U., Pickard, J.M., Gillilland, M.G., Pudlo, N.A., Hoostal, M., Li, X., Wang, T.D., Feehley, T., et al. (2017). Neonatal acquisition of Clostridia species protects against colonization by bacterial pathogens. Science (1979). 356, 315–319.

25. Sivan, A., Corrales, L., Hubert, N., Williams, J.B., Aquino-Michaels, K., Earley, Z.M., Benyamin, F.W., Lei, Y.M., Jabri, B., Alegre, M.-L., et al. (2015). Commensal Bifidobacterium promotes antitumor immunity and facilitates anti–PD-L1 efficacy. Science (1979). 350, 1084–1089. 10.1126/science.aad1329.

26. Littmann, E.R., Lee, J.J., Denny, J.E., Alam, Z., Maslanka, J.R., Zarin, I., Matsuda, R., Carter, R.A., Susac, B., Saffern, M.S., et al. (2021). Host immunity modulates the efficacy of microbiota transplantation for treatment of Clostridioides difficile infection. Nat. Commun. 12, 1–15. 10.1038/s41467-020-20793-x.

27. Binion, D.G. (2012). Clostridium difficile Infection in Patients with Inflammatory Bowel Disease.

28. Nguyen, G.C., Kaplan, G.G., Harris, M.L., and Brant, S.R. (2008). A national survey of the prevalence and impact of Clostridium difficile infection among hospitalized inflammatory bowel disease patients. Preprint, 10.1111/j.1572-0241.2007.01780.x https://doi.org/10.1111/j.1572-0241.2007.01780.x.

29. Hourigan, S.K., Oliva-Hemker, M., and Hutfless, S. (2014). The Prevalence of Clostridium difficile Infection in Pediatric and Adult Patients with Inflammatory Bowel Disease. Dig. Dis. Sci. 59, 2222–2227. 10.1007/s10620-014-3169-4.

30. Ananthakrishnan, A.N. (2012). Detecting and Treating Clostridium Difficile Infections in Patients with Inflammatory Bowel Disease. Preprint, 10.1016/j.gtc.2012.01.003 https://doi.org/10.1016/j.gtc.2012.01.003.

31. Qi, H.-X., Wang, Q., and Zhou, G.-Q. (2025). Association of Clostridium difficile infection with clinical outcomes of patients with inflammatory bowel disease: A meta-analysis . World J. Gastrointest. Surg. 17. 10.4240/wjgs.v17.i4.100555.

32. Strober, W., Fuss, I., and Mannon, P. (2007). The fundamental basis of inflammatory bowel disease. Journal of Clinical Investigation 117, 514–521. 10.1172/JCI30587.

33. Ramos, G.P., and Papadakis, K.A. (2019). Mechanisms of Disease: Inflammatory Bowel Diseases. Preprint at Elsevier Ltd, 10.1016/j.mayocp.2018.09.013 https://doi.org/10.1016/j.mayocp.2018.09.013.

34. Hvas, C.L., Dahl Jørgensen, S.M., Jørgensen, S.P., Storgaard, M., Lemming, L., Hansen, M.M., Erikstrup, C., and Dahlerup, J.F. (2019). Fecal Microbiota Transplantation Is Superior to Fidaxomicin for Treatment of Recurrent Clostridium difficile Infection. Gastroenterology 156, 1324–1332.e3. 10.1053/j.gastro.2018.12.019.

35. Minkoff, N.Z., Aslam, S., Medina, M., Tanner-Smith, E.E., Zackular, J.P., Acra, S., Nicholson, M.R., and Imdad, A. (2023). Fecal microbiota transplantation for the treatment of recurrent Clostridioides difficile (Clostridium difficile). Cochrane Database of Systematic Reviews 2023. 10.1002/14651858.CD013871.pub2.

36. Khanna, S., Vazquez-Baeza, Y., González, A., Weiss, S., Schmidt, B., Muñiz-Pedrogo, D.A., Rainey, J.F., Kammer, P., Nelson, H., Sadowsky, M., et al. (2017). Changes in microbial ecology after fecal microbiota transplantation for recurrent C. difficile infection affected by underlying inflammatory bowel disease. Microbiome 5. 10.1186/S40168-017-0269-3.

37. Fischer, M., Kao, D., Mehta, S.R., Martin, T., Dimitry, J., Keshteli, A.H., Cook, G.K., Phelps, E., Sipe, B.W., Xu, H., et al. (2016). Predictors of Early Failure after Fecal Microbiota Transplantation for the Therapy of Clostridium Difficile Infection: A Multicenter Study. American Journal of Gastroenterology 111, 1024–1031. 10.1038/ajg.2016.180.

38. Khoruts, A., Rank, K.M., Newman, K.M., Viskocil, K., Vaughn, B.P., Hamilton, M.J., and Sadowsky, M.J. (2016). Inflammatory Bowel Disease Affects the Outcome of Fecal Microbiota Transplantation for Recurrent Clostridium difficile Infection. Clinical Gastroenterology and Hepatology 14, 1433–1438. 10.1016/j.cgh.2016.02.018.

39. Kelsen, J.R., Kim, J., Latta, D., Smathers, S., McGowan, K.L., Zaoutis, T., Mamula, P., and Baldassano, R.N. (2011). Recurrence rate of clostridium difficile infection in hospitalized pediatric patients with inflammatory bowel disease. Inflamm. Bowel Dis. 17, 50–55. 10.1002/ibd.21421.

40. Orth, P., Xiao, L., Hernandez, L.D., Reichert, P., Sheth, P.R., Beaumont, M., Yang, X., Murgolo, N., Ermakov, G., Dinunzio, E., et al. (2014). Mechanism of action and epitopes of Clostridium difficile toxin B-neutralizing antibody bezlotoxumab revealed by X-ray crystallography. Journal of Biological Chemistry 289, 18008–18021. 10.1074/jbc.M114.560748.

41. Wilcox, M.H., Gerding, D.N., Poxton, I.R., Kelly, C., Nathan, R., Birch, T., Cornely, O.A., Rahav, G., Bouza, E., Lee, C., et al. (2017). Bezlotoxumab for Preventing Recurrent Clostridium difficile Infections. New England Journal of Medicine 376, 305–317. 10.1056/nejmoa1602615.

42. Fein, A., Kern, C., Barrett, T., and Perry, C. (2022). Bezlotoxumab Therapy for Recurrent Clostridium difficile Infection in an Ulcerative Colitis Patient. Crohns Colitis 360 4. 10.1093/crocol/otac038.

43. Singh, P. (2024). Merck to discontinue drug for bacterial infection. Reuters.

44. Cribas, E.S., Denny, J.E., Maslanka, J.R., and Abt, M.C. (2021). Loss of interleukin-10 (IL-10) signaling promotes IL-22Dependent host defenses against acute clostridioides difficile infection. Infect. Immun. 89, 22. 10.1128/IAI.00730-20.

45. Saraiva, M., and O’Garra, A. (2010). The regulation of IL-10 production by immune cells. Preprint, 10.1038/nri2711 https://doi.org/10.1038/nri2711.

46. Strober, W., Fuss, I.J., and Blumberg, R.S. (2002). The immunology of mucosal models of inflammation. Preprint, 10.1146/annurev.immunol.20.100301.064816 https://doi.org/10.1146/annurev.immunol.20.100301.064816.

47. K∼jhn, R., L6hler, I., Rennick, D., Rajewsky, K., and Moiler, W. (1993). Interleukin-lO-Deficient Mice Develop Chronic Enterocolitis.

48. Seekatz, A.M., Aas, J., Gessert, C.E., Rubin, T.A., Saman, D.M., Bakken, J.S., and Young, V.B. (2014). Recovery of the gut microbiome following fecal microbiota transplantation. mBio 5. 10.1128/mBio.00893-14.

49. Weingarden, A.R., Chen, C., Bobr, A., Yao, D., Lu, Y., Nelson, V.M., Sadowsky, M.J., and Khoruts, A. (2014). Microbiota transplantation restores normal fecal bile acid composition in recurrent Clostridium difficile infection. J Physiol Gastrointest Liver Physiol 306, 310–319. 10.1152/ajpgi.00282.2013.-Fecal.

50. Weingarden, A.R., Dosa, P.I., DeWinter, E., Steer, C.J., Shaughnessy, M.K., Johnson, J.R., Khoruts, A., and Sadowsky, M.J. (2016). Changes in colonic bile acid composition following fecal microbiota transplantation are sufficient to control Clostridium difficile germination and growth. PLoS One 11. 10.1371/journal.pone.0147210.

51. Seekatz, A.M., Theriot, C.M., Rao, K., Chang, Y.M., Freeman, A.E., Kao, J.Y., and Young, V.B. (2018). Restoration of short chain fatty acid and bile acid metabolism following fecal microbiota transplantation in patients with recurrent Clostridium difficile infection. Anaerobe 53, 64–73. 10.1016/j.anaerobe.2018.04.001.

52. Lochner, M., Peduto, L., Cherrier, M., Sawa, S., Langa, F., Varona, R., Riethmacher, D., Si-Tahar, M., Di Santo, J.P., and Eberl, G. (2008). In vivo equilibrium of proinflammatory IL-17+ and regulatory IL-10+ Foxp3+ RORγt+ T cells. Journal of Experimental Medicine 205, 1381–1393. 10.1084/jem.20080034.

53. Koch, M.A., Tucker-Heard, G., Perdue, N.R., Killebrew, J.R., Urdahl, K.B., and Campbell, D.J. (2009). The transcription factor T-bet controls regulatory T cell homeostasis and function during type 1 inflammation. Nat. Immunol. 10, 595–602. 10.1038/ni.1731.

54. Yadav, M., Stephan, S., and Bluestone, J.A. (2013). Peripherally induced Tregs-role in immune homeostasis and autoimmunity. Front. Immunol. 4. 10.3389/fimmu.2013.00232.

55. Cheru, N., Hafler, D.A., and Sumida, T.S. (2023). Regulatory T cells in peripheral tissue tolerance and diseases. Preprint at Frontiers Media S.A., 10.3389/fimmu.2023.1154575 https://doi.org/10.3389/fimmu.2023.1154575.

56. Wing, K., Onishi, Y., Prieto-Martin, P., Yamaguchi, T., Miyara, M., Fehervari, Z., Nomura, T., and Sakaguchi, S. (2008). CTLA-4 Control over Foxp3+ Regulatory T Cell Function. Science (1979). 322, 271–275. 10.1126/science.1164164.

57. Wing, J.B., Ise, W., Kurosaki, T., and Sakaguchi, S. (2014). Regulatory T cells control antigen-specific expansion of Tfh cell number and humoral immune responses via the coreceptor CTLA-4. Immunity 41, 1013–1025. 10.1016/j.immuni.2014.12.006.

58. Sage, P.T., Paterson, A.M., Lovitch, S.B., and Sharpe, A.H. (2014). The coinhibitory receptor CTLA-4 controls B cell responses by modulating T follicular helper, T follicular regulatory, and T regulatory cells. Immunity 41, 1026–1039. 10.1016/j.immuni.2014.12.005.

59. Shafiani, S., Dinh, C., Ertelt, J.M., Moguche, A.O., Siddiqui, I., Smigiel, K.S., Sharma, P., Campbell, D.J., Way, S.S., and Urdahl, K.B. (2013). Pathogen-Specific Treg Cells Expand Early during Mycobacterium tuberculosis Infection but Are Later Eliminated in Response to Interleukin-12. Immunity 38, 1261–1270. 10.1016/j.immuni.2013.06.003.

60. Perry, J.A., Shallberg, L., Clark, J.T., Gullicksrud, J.A., DeLong, J.H., Douglas, B.B., Hart, A.P., Lanzar, Z., O’Dea, K., Konradt, C., et al. (2022). PD-L1–PD-1 interactions limit effector regulatory T cell populations at homeostasis and during infection. Nat. Immunol. 23, 743–756. 10.1038/s41590-022-01170-w.

61. Pereira, J.A., Lanzar, Z., Clark, J.T., Hart, A.P., Douglas, B.B., Shallberg, L., O’Dea, K., Christian, D.A., and Hunter, C.A. (2023). PD-1 and CTLA-4 exert additive control of effector regulatory T cells at homeostasis. Front. Immunol. 14. 10.3389/fimmu.2023.997376.

62. Li, D.Y., and Xiong, X.Z. (2020). ICOS+ Tregs: A Functional Subset of Tregs in Immune Diseases. Preprint at Frontiers Media S.A., 10.3389/fimmu.2020.02104 https://doi.org/10.3389/fimmu.2020.02104.

63. Li, D.Y., and Xiong, X.Z. (2020). ICOS+ Tregs: A Functional Subset of Tregs in Immune Diseases. Preprint at Frontiers Media S.A., 10.3389/fimmu.2020.02104 https://doi.org/10.3389/fimmu.2020.02104.

64. Bain, C.C., Scott, C.L., Uronen-Hansson, H., Gudjonsson, S., Jansson, O., Grip, O., Guilliams, M., Malissen, B., Agace, W.W., and Mowat, A.M.I. (2013). Resident and pro-inflammatory macrophages in the colon represent alternative context-dependent fates of the same Ly6C hi monocyte precursors. Mucosal Immunol. 6, 498–510. 10.1038/mi.2012.89.

65. Kayama, H., Ueda, Y., Sawa, Y., Jeon, S.G., Ma, J.S., Okumura, R., Kubo, A., Ishii, M., Okazaki, T., Murakami, M., et al. (2012). Intestinal CX 3C chemokine receptor 1 high (CX 3CR1 high) myeloid cells prevent T-cell-dependent colitis. Proc. Natl. Acad. Sci. U. S. A. 109, 5010–5015. 10.1073/pnas.1114931109.

66. Rivera-Chávez, F., Zhang, L.F., Faber, F., Lopez, C.A., Byndloss, M.X., Olsan, E.E., Xu, G., Velazquez, E.M., Lebrilla, C.B., Winter, S.E., et al. (2016). Depletion of Butyrate-Producing Clostridia from the Gut Microbiota Drives an Aerobic Luminal Expansion of Salmonella. Cell Host Microbe 19, 443–454. 10.1016/j.chom.2016.03.004.

67. Litvak, Y., Mon, K.K.Z., Nguyen, H., Chanthavixay, G., Liou, M., Velazquez, E.M., Kutter, L., Alcantara, M.A., Byndloss, M.X., Tiffany, C.R., et al. (2019). Commensal Enterobacteriaceae Protect against Salmonella Colonization through Oxygen Competition. Cell Host Microbe 25, 128–139.e5. 10.1016/j.chom.2018.12.003.

68. Shelton, C.D., Yoo, W., Shealy, N.G., Torres, T.P., Zieba, J.K., Calcutt, M.W., Foegeding, N.J., Kim, D., Kim, J., Ryu, S., et al. (2022). Salmonella enterica serovar Typhimurium uses anaerobic respiration to overcome propionate-mediated colonization resistance. Cell Rep. 38. 10.1016/j.celrep.2021.110180.

69. Thiennimitr, P., Winter, S.E., Winter, M.G., Xavier, M.N., Tolstikov, V., Huseby, D.L., Sterzenbach, T., Tsolis, R.M., Roth, J.R., and Bäumler, A.J. (2011). Intestinal inflammation allows Salmonella to use ethanolamine to compete with the microbiota. Proc. Natl. Acad. Sci. U. S. A. 108, 17480–17485. 10.1073/pnas.1107857108.

70. Lopez, C.A., Miller, B.M., Rivera-Chávez, F., Velazquez, E.M., Byndloss, M.X., Chávez-Arroyo, A., Lokken, K.L., Tsolis, R.M., Winter, S.E., and Bäumler, A.J. (2016). Virulence factors enhance Citrobacter rodentium expansion through aerobic respiration.

71. Byndloss, M.X., Olsan, E.E., Rivera-Chávez, F., Tiffany, C.R., Cevallos, S.A., Lokken, K.L., Torres, T.P., Byndloss, A.J., Faber, F., Gao, Y., et al. (2017). Microbiota-activated PPAR-g signaling inhibits dysbiotic Enterobacteriaceae expansion.

72. Winter, S.E., Thiennimitr, P., Winter, M.G., Butler, B.P., Huseby, D.L., Crawford, R.W., Russell, J.M., Bevins, C.L., Adams, L.G., Tsolis, R.M., et al. (2010). Gut inflammation provides a respiratory electron acceptor for Salmonella. Nature 467, 426–429. 10.1038/nature09415.

73. Winter, S.E., Winter, M.G., Xavier, M.N., Thiennimitr, P., Poon, V., Keestra, A.M., Laughlin, R.C., Gomez, G., Wu, J., Lawhon, S.D., et al. (2013). Host-Derived Nitrate Boosts Growth of E. coli in the Inflamed Gut.

74. Wang, S., El-Fahmawi, A., Christian, D.A., Fang, Q., Radaelli, E., Chen, L., Sullivan, M.C., Misic, A.M., Ellringer, J.A., Zhu, X.-Q., et al. (2019). Infection-Induced Intestinal Dysbiosis Is Mediated by Macrophage Activation and Nitrate Production. 10.1128/mBio.

75. Winterbourn, C.C., Kettle, A.J., and Hampton, M.B. (2016). Reactive Oxygen Species and Neutrophil Function. Annu. Rev. Biochem. 85, 765–792. 10.1146/annurev-biochem-060815-014442.

76. Huang, X., Li, J., Dorta-Estremera, S., Di Domizio, J., Anthony, S.M., Watowich, S.S., Popkin, D., Liu, Z., Brohawn, P., Yao, Y., et al. (2015). Neutrophils Regulate Humoral Autoimmunity by Restricting Interferon-γ Production via the Generation of Reactive Oxygen Species. Cell Rep. 12, 1120–1132. 10.1016/j.celrep.2015.07.021.

77. Veenith, T., Martin, H., Le Breuilly, M., Whitehouse, T., Gao-Smith, F., Duggal, N., Lord, J.M., Mian, R., Sarphie, D., and Moss, P. (2022). High generation of reactive oxygen species from neutrophils in patients with severe COVID-19. Sci. Rep. 12. 10.1038/s41598-022-13825-7.

78. Fialkow, L., Wang, Y., and Downey, G.P. (2007). Reactive oxygen and nitrogen species as signaling molecules regulating neutrophil function. Preprint, 10.1016/j.freeradbiomed.2006.09.030 https://doi.org/10.1016/j.freeradbiomed.2006.09.030.

79. Wang, S., El-Fahmawi, A., Christian, D.A., Fang, Q., Radaelli, E., Chen, L., Sullivan, M.C., Misic, A.M., Ellringer, J.A., Zhu, X.-Q., et al. (2019). Infection-Induced Intestinal Dysbiosis Is Mediated by Macrophage Activation and Nitrate Production. 10.1128/mBio.

80. Winter, S.E., Winter, M.G., Xavier, M.N., Thiennimitr, P., Poon, V., Keestra, A.M., Laughlin, R.C., Gomez, G., Wu, J., Lawhon, S.D., et al. (2013). Host-derived nitrate boosts growth of E. coli in the inflamed gut. Science (1979). 339, 708–711. 10.1126/science.1232467.

81. Wang, C., Chen, K., Xia, Y., Dai, W., Wang, F., Shen, M., Cheng, P., Wang, J., Lu, J., Zhang, Y., et al. (2014). N-Acetylcysteine attenuates ischemia-reperfusion-induced apoptosis and autophagy in mouse liver via regulation of the ROS/JNK/Bcl-2 Pathway. PLoS One 9. 10.1371/journal.pone.0108855.

82. Halasi, M., Wang, M., Chavan, T.S., Gaponenko, V., Hay, N., and Gartel, A.L. (2013). ROS inhibitor N-acetyl-L-cysteine antagonizes the activity of proteasome inhibitors. Biochemical Journal 454, 201–208. 10.1042/BJ20130282.

83. Gesser, B., Leffers, H., Jinquan, T., Vestergaard, C., Kirstein, N., Sindet-Pedersen, S., Lindkaer Jensen, S., Thestrup-Pedersen, K., and Larsen, C.G. (1997). Identification of functional domains on human interleukin 10.

84. Thompson-Snipes, L., Dhar, V., Bond, M.W., Mosmann, T.R., Moore, K.W., and Rennick, D.M. Interleukin 10: A Novel Stimulatory Factor for Mast Cells and Their Progenitors.

85. Wei, Y.L., Chen, Y.Q., Gong, H., Li, N., Wu, K.Q., Hu, W., Wang, B., Liu, K.J., Wen, L.Z., Xiao, X., et al. (2018). Fecal microbiota transplantation ameliorates experimentally induced colitis in mice by upregulating AhR. Front. Microbiol. 9. 10.3389/fmicb.2018.01921.

86. Burrello, C., Garavaglia, F., Cribiù, F.M., Ercoli, G., Lopez, G., Troisi, J., Colucci, A., Guglietta, S., Carloni, S., Guglielmetti, S., et al. (2018). Therapeutic faecal microbiota transplantation controls intestinal inflammation through IL10 secretion by immune cells. Nat. Commun. 9. 10.1038/s41467-018-07359-8.

87. Danne, C., Rolhion, N., and Sokol, H. (2021). Recipient factors in faecal microbiota transplantation: one stool does not fit all. Preprint at Nature Research, 10.1038/s41575-021-00441-5 https://doi.org/10.1038/s41575-021-00441-5.

88. Steidler, L., Hans, W., Schotte, L., Neirynck, S., Obermeier, F., Falk, W., Fiers, W., and Remaut, E. (1994). Treatment of Murine Colitis by Lactococcus lactis Secreting Interleukin-10.

89. Duchmann, R., Schmitt, E., Knolle, P., Zum Büschenfelde, K.H.M., and Neurath, M. (1996). Tolerance towards resident intestinal flora in mice is abrogated in experimental colitis and restored by treatment with interleukin-10 or antibodies to interleukin-12. Eur. J. Immunol. 26, 934–938. 10.1002/eji.1830260432.

90. Lindsay, J.O., Ciesielski, C.J., Scheinin, T., Hodgson, H.J., and Brennan, F.M. (2001). The Prevention and Treatment of Murine Colitis Using Gene Therapy with Adenoviral Vectors Encoding IL-101. The Journal of Immunology 166, 7625–7633.

91. Zurita-Turk, M., del Carmen, S., Santos, A.C.G., Pereira, V.B., Cara, D.C., Leclercq, S.Y., de LeBlanc, A.D.M., Azevedo, V., Chatel, J.M., LeBlanc, J.G., et al. (2014). Lactococcus lactis carrying the pValac DNA expression vector coding for IL-10 reduces inflammation in a murine model of experimental colitis. BMC Biotechnol. 14. 10.1186/1472-6750-14-73.

92. Marlow, G.J., van Gent, D., and Ferguson, L.R. (2013). Why interleukin-10 supplementation does not work in Crohn’s disease patients. World J. Gastroenterol. 19, 3931–3941. 10.3748/wjg.v19.i25.3931.

93. Fedorak, R.N., Gangl, A., Elson, C.O., Rutgeerts, P., Schreiber, S., Wild, G., Hanauer, S.B., Kilian, A., Cohard, M., LeBeaut, A., et al. (2000). Recombinant human interleukin 10 in the treatment of patients with mild to moderately active Crohn’s disease. Gastroenterology 119, 1473–1482. 10.1053/gast.2000.20229.

94. Aunins, E.A., Phan, A.T., Alameh, M.-G., Dwivedi, G., Cruz-Morales, E., Christian, D.A., Tam, Y., Bunkofske, M.E., Zabala Peñafiel, A., O, K.M., et al. (2025). An Il12 mRNA-LNP adjuvant enhances mRNA vaccine-induced CD8 T cell responses.

95. Sung, J., Alghoul, Z., Long, D., Yang, C., and Merlin, D. (2022). Oral delivery of IL-22 mRNA-loaded lipid nanoparticles targeting the injured intestinal mucosa: A novel therapeutic solution to treat ulcerative colitis. Biomaterials 288. 10.1016/j.biomaterials.2022.121707.

96. Gál, L., Bellák, T., Marton, A., Fekécs, Z., Weissman, D., Török, D., Biju, R., Vizler, C., Kristóf, R., Beattie, M.B., et al. (2023). Restoration of Motor Function through Delayed Intraspinal Delivery of Human IL-10-Encoding Nucleoside-Modified mRNA after Spinal Cord Injury. Research 6. 10.34133/research.0056.

97. Liu, C., Huang, X., Chen, K.S., Xiong, S., Yaremenko, A. V., Zhen, X., You, X., Rossignoli, F., Tang, Y., Koo, S., et al. (2025). Systemic reprogramming of tumour immunity via IL-10-mRNA nanoparticles. Nat. Nanotechnol. 20, 1526–1538. 10.1038/s41565-025-01980-7.

98. Reyes-Esteves, S., Majumder, A., Marzolini, N., Zamora, M.E., Jeong, S., Wang, Y., Patel, M.N., Espy, C., Papp, T.E., Akyianu, A., et al. (2025). Targeted lipid nanoparticles containing IL-10 mRNA improve outcomes in experimental intracerebral hemorrhage. Journal of Neuroinflammation 22. 10.1186/s12974-025-03541-0.

99. Veiga, N., Goldsmith, M., Granot, Y., Rosenblum, D., Dammes, N., Kedmi, R., Ramishetti, S., and Peer, D. (2018). Cell specific delivery of modified mRNA expressing therapeutic proteins to leukocytes. Nat. Commun. 9. 10.1038/s41467-018-06936-1.

100. Monday, L., Tillotson, G., and Chopra, T. (2024). Microbiota-Based Live Biotherapeutic Products for Clostridioides Difficile Infection-The Devil is in the Details. Preprint at Dove Medical Press Ltd, 10.2147/IDR.S419243 https://doi.org/10.2147/IDR.S419243.

101. Khanna, S., Pardi, D.S., Jones, C., Shannon, W.D., Gonzalez, C., and Blount, K. (2021). RBX7455, a Non-frozen, Orally Administered Investigational Live Biotherapeutic, Is Safe, Effective, and Shifts Patients’ Microbiomes in a Phase 1 Study for Recurrent Clostridioides difficile Infections. Clinical Infectious Diseases 73, E1613–E1620. 10.1093/cid/ciaa1430.

102. Allegretti, J.R., Kelly, C.R., Louie, T., Fischer, M., Hota, S., Misra, B., Van Hise, N.W., Yen, E., Bullock, J.S., Silverman, M., et al. (2025). Safety and Tolerability of CP101, a Full-Spectrum, Oral Microbiome Therapeutic for the Prevention of Recurrent Clostridioides difficile Infection: A Phase 2 Randomized Controlled Trial. Gastroenterology 168, 357–366.e3. 10.1053/j.gastro.2024.09.030.

103. Lee, C., Louie, T., Bancke, L., Guthmueller, B., Harvey, A., Feuerstadt, P., Khanna, S., Orenstein, R., and Dubberke, E.R. (2023). Safety of fecal microbiota, live-jslm (REBYOTA^TM^) in individuals with recurrent Clostridioides difficile infection: data from five prospective clinical trials. Therap. Adv. Gastroenterol. 16. 10.1177/17562848231174277.

104. Khanna, S., Pardi, D.S., Kelly, C.R., Kraft, C.S., Dhere, T., Henn, M.R., Lombardo, M.J., Vulic, M., Ohsumi, T., Winkler, J., et al. (2016). A Novel Microbiome Therapeutic Increases Gut Microbial Diversity and Prevents Recurrent Clostridium difficile Infection. Journal of Infectious Diseases 214, 173–181. 10.1093/infdis/jiv766.

105. Wang, Y., Hunt, A., Danziger, L., and Drwiega, E.N. (2024). A Comparison of Currently Available and Investigational Fecal Microbiota Transplant Products for Recurrent Clostridioides difficile Infection. Preprint at Multidisciplinary Digital Publishing Institute (MDPI), 10.3390/antibiotics13050436 https://doi.org/10.3390/antibiotics13050436.

106. Stallhofer, J., Steube, A., Katzer, K., and Stallmach, A. (2024). Microbiota-Based Therapeutics as New Standard-of-Care Treatment for Recurrent Clostridioides difficile Infection. Preprint at S. Karger AG, 10.1159/000535851 https://doi.org/10.1159/000535851.

107. Maslanka, J.R., Gu, C.H., Zarin, I., Denny, J.E., Broadaway, S., Fett, B., Mattei, L.M., Walk, S.T., and Abt, M.C. (2020). Detection and elimination of a novel non-toxigenic Clostridioides difficile strain from the microbiota of a mouse colony. Gut Microbes 12, 1–15. 10.1080/19490976.2020.1851999.

108. Fu Wang, R., and Kushner, S.R. (1991). Construction of versatile low-copy-number vectors for cloning, sequencing and gene expression in Escherichia coli (Recombinant DNA; polymerase chain reaction; deletion analysis; ampicillin resistance; single-stranded DNA; kanamycin resistance).

109. Denny, J.E., Flores, J.N., Mdluli, N. V., and Abt, M.C. (2025). Standard mouse diets lead to differences in severity in infectious and non-infectious colitis. mBio 16. 10.1128/mbio.03302-24.

110. Guerrini, C.J., Botkin, J.R., and McGuire, A.L. (2019). Reproducible, interactive, scalable and extensible microbiome data science using QIIME 2. Preprint at Nature Publishing Group, 10.1038/s41587-019-0190-3 https://doi.org/10.1038/s41587-019-0190-3.

111. Callahan, B.J., McMurdie, P.J., Rosen, M.J., Han, A.W., Johnson, A.J.A., and Holmes, S.P. (2016). DADA2: High-resolution sample inference from Illumina amplicon data. Nat. Methods 13, 581–583. 10.1038/nmeth.3869.

112. Bokulich, N.A., Kaehler, B.D., Rideout, J.R., Dillon, M., Bolyen, E., Knight, R., Huttley, G.A., and Gregory Caporaso, J. (2018). Optimizing taxonomic classification of marker-gene amplicon sequences with QIIME 2’s q2-feature-classifier plugin. Microbiome 6. 10.1186/s40168-018-0470-z.

113. McDonald, D., Jiang, Y., Balaban, M., Cantrell, K., Zhu, Q., Gonzalez, A., Morton, J.T., Nicolaou, G., Parks, D.H., Karst, S.M., et al. (2024). Greengenes2 unifies microbial data in a single reference tree. Nat. Biotechnol. 42, 715–718. 10.1038/s41587-023-01845-1.

114. R Core Team (2023). _R: A Language and Environment for Statistical Computing_. R Foundation for Statistical Computing, Vienna, Austria. https://www.R-project.org/. Preprint.

115. McMurdie, P.J., and Holmes, S. (2013). Phyloseq: An R Package for Reproducible Interactive Analysis and Graphics of Microbiome Census Data. PLoS One 8. 10.1371/journal.pone.0061217.

116. Create Elegant Data Visualisations Using the Grammar of Graphics.

117. Lozupone, C., and Knight, R. (2005). UniFrac: A new phylogenetic method for comparing microbial communities. Appl. Environ. Microbiol. 71, 8228–8235. 10.1128/AEM.71.12.8228-8235.2005.

118. Lachenbruch, P.A. (2001). Comparisons of two-part models with competitors. Stat. Med. 20, 1215–1234. 10.1002/sim.790.

119. Lachenbruch, P.A. (2001). Power and sample size requirements for two-part models. Stat. Med. 20, 1235–1238. 10.1002/sim.812.

120. Package “vegan” Title Community Ecology Package (2025).

