## Supplementary material for "Immunotherapy restores therapeutic efficacy of fecal microbiome transplants to treat *Clostridioides difficile*": Supplmental Tables

| Pair comparison | $R^2$ | p-value | Adj p-value (FDR) |
| --- | --- | --- | --- |
| **Fig1B (Weighted UniFrac distance)** |  |  |  |
| IL10-/- vs IL10HET post-FMT D0 | 0.2838531 | 0.129 | 0.129 |
| IL10-/- vs IL10HET post-FMT D8 | 0.57423317 | 0.021 | 0.042 |
| **FigS1C (Unweighted UniFrac distance)** |  |  |  |
| IL10-/- vs IL10HET post-FMT D0 | 0.46654491 | 0.061 | 0.061 |
| IL10-/- vs IL10HET post-FMT D8 | 0.41364811 | 0.036 | 0.061 |
| **Fig1E (Weighted UniFrac distance)** |  |  |  |
| B6 + algG vs B6 + aIL10R post-FMT D0 | 0.09605401 | 0.651 | 0.651 |
| B6 + algG vs B6 + aIL10R post-FMT D8 | 0.59226665 | 0.029 | 0.058 |
| **FigS1E (Weighted UniFrac distance)** |  |  |  |
| ABX_IL10HET vs ABX_IL10-/- post-FMT D0 | 0.07335485 | 0.737 | 0.737 |
| ABX_IL10HET vs ABX_IL10-/- post-FMT D4 | 0.47317621 | 0.066 | 0.099 |
| ABX_IL10HET vs ABX_IL10-/- post-FMT D12 | 0.57250581 | 0.016 | 0.048 |
| **FigS1G (Unweighted UniFrac distance)** |  |  |  |
| B6 + algG vs B6 + aIL10R post-FMT D0 | 0.06442526 | 0.818 | 0.818 |
| B6 + algG vs B6 + aIL10R post-FMT D8 | 0.34845096 | 0.038 | 0.076 |
| **Fig5E (Weighted UniFrac distance)** |  |  |  |
| Foxp3DTR + PBS vs DT_Il10_LNP post-FMT D0 | 0.18221321 | 0.24 | 0.324 |
| Foxp3DTR + PBS vs DT_empty_LNP post-FMT D0 | 0.10680529 | 0.648 | 0.648 |
| Foxp3DTR + DT_IL10_LNP vs DT_empty_LNP post-FMT D0 | 0.12573897 | 0.27 | 0.324 |
| Foxp3DTR + PBS vs DT_Il10_LNP post-FMT D8 | 0.45510988 | 0.012 | 0.042 |
| Foxp3DTR + PBS vs DT_empty_LNP post-FMT D8 | 0.64679753 | 0.058 | 0.116 |
| Foxp3DTR + DT_IL10_LNP vs DT_empty_LNP post-FMT D8 | 0.36770238 | 0.014 | 0.042 |
| **FigS5F (Unweighted UniFrac distance)** |  |  |  |
| Foxp3DTR + PBS vs DT_Il10_LNP post-FMT D0 | 0.07538627 | 0.849 | 0.981 |
| Foxp3DTR + PBS vs DT_empty_LNP post-FMT D0 | 0.07409969 | 0.981 | 0.981 |
| Foxp3DTR + DT_IL10_LNP vs DT_empty_LNP post-FMT D0 | 0.08112592 | 0.763 | 0.981 |
| Foxp3DTR + PBS vs DT_Il10_LNP post-FMT D8 | 0.24544011 | 0.012 | 0.045 |
| Foxp3DTR + PBS vs DT_empty_LNP post-FMT D8 | 0.48043651 | 0.055 | 0.11 |
| Foxp3DTR + DT_IL10_LNP vs DT_empty_LNP post-FMT D8 | 0.30222712 | 0.015 | 0.045 |

**Supplementary Table 1:** PERMANOVA analysis of weighted and unweighted UniFrac distances between the intestinal microbial communities for experimental groups in different experiments shown in respective figures. PERMANOVA test tested the null hypothesis that there is no difference between the microbial compositions of test groups. The Benjamini-Hochberg method was used to adjust for multiple comparisons.

|  | Infected | Resolved | Total mice | Percentage resolved |
| --- | --- | --- | --- | --- |
| **Data from Fig1A** |  |  |  |  |
| IL10HET | 3 | 11 | 14 | 78.57 |
| IL10-/- | 9 | 1 | 10 | 10.00 |
| **Data from Fig1D** |  |  |  |  |
| B6 + α-IL10RB | 9 | 2 | 11 | 18.18 |
| B6 + Isotype | 1 | 8 | 9 | 88.89 |
| **Data from FigS1J** |  |  |  |  |
| B6 + IL10RB-D7 | 6 | 2 | 8 | 25.00 |
| B6 + IL10RB-D21 | 4 | 4 | 8 | 50.00 |
| B6 + Isotype | 2 | 5 | 7 | 71.43 |
| **Data from Fig2E** |  |  |  |  |
| CD4_Cre+: IL10 F/F | 5 | 1 | 6 | 16.67 |
| Cd4_Cre+ | 0 | 7 | 7 | 100.00 |
| **Data from Fig3Q** |  |  |  |  |
| Foxp3DTR + DT + Iso-FMT | 7 | 0 | 7 | 0.00 |
| Foxp3DTR + DT + Anti-IFNγ-FMT | 2 | 6 | 8 | 75.00 |
| Foxp3DTR + PBS + Iso-FMT | 0 | 7 | 7 | 100.00 |
| Foxp3DTR + PBS + Iso-noFMT | 7 | 0 | 7 | 0.00 |
| **Data from Fig4C** |  |  |  |  |
| Foxp3DTR + PBS + Isotype | 2 | 8 | 10 | 80.00 |
| Foxp3DTR + DT + Isotype | 9 | 2 | 11 | 18.18 |
| Foxp3DTR + DT + αLy6g | 7 | 5 | 12 | 41.67 |
| **Data from Fig4D** |  |  |  |  |
| Foxp3DTR + PBS | 0 | 5 | 5 | 100.00 |
| Foxp3DTR + DT | 2 | 2 | 4 | 50.00 |
| Foxp3DTR + DT + AG + NAC | 1 | 6 | 7 | 85.71 |
| **Data from Fig4E** |  |  |  |  |
| B6 + Isotype | 3 | 7 | 10 | 70.00 |
| B6 + IL10R | 9 | 1 | 10 | 10.00 |
| B6 + IL10R + AG + NAC | 7 | 6 | 13 | 46.15 |
| **Data from Fig5C** |  |  |  |  |
| Foxp3DTR + PBS | 0 | 3 | 3 | 100.00 |
| Foxp3DTR + DT-mRNA(empty) | 3 | 1 | 4 | 25.00 |
| Foxp3DTR + DT-mRNA(il10) | 1 | 5 | 6 | 83.33 |

**Supplementary Table 2:** Percentage of mice that resolved *C. difficile* infection at Day 12 post-FMT. Number of infected vs resolved, and the total number of mice used in each experimental

group for different experiments shown in respective figures. For *Il10*-mRNA-LNP experiment. Data from Fig 5C is Day 11 post-FMT.

| **Marker** | **Clone** | **Fluorochrome** | **Catalog#** | **Dilution** | **Company** |
| --- | --- | --- | --- | --- | --- |
| Ly6g | 1A8 | A700 | 127622 | 100 | Biolegend |
| CD45 | 30-F11 | A700 | 103128 | 100 | Biolegend |
| IFN-g | XMG1.2 | APC | 17-7311-82 | 200 | Invitrogen |
| CD4 | RM4-5 | BV605 | 100548 | 200 | Biolegend |
| CD45 | 30-F11 | BV605 | 103155 | 200 | Biolegend |
| CD19 | 6D5 | BV785 | 115543 | 200 | Biolegend |
| CD90.2 | 53-2.1 | eF450 | 48-0902-82 | 300 | eBioscience |
| MHC-II (I-Ab) | AF6-120.1 | eF450 | 48-5320-82 | 200 | eBioscience |
| Ly6c | HK1.4 | eF780 | 47-5932-82 | 200 | eBioscience |
| IL17A | TC11-18H10.1 | FITC | 506908 | 300 | Biolegend |
| Gr-1 | RB6-8C5 | PE Dazzle 594 | 108452 | 300 | Biolegend |
| CD3e | 1452C11 | PerCP Cy5.5 | 45-0031-82 | 200 | eBioscience |
| CD5 | 53-7.3 | PerCp Cy5.5 | 45-0051-82 | 300 | eBioscience |
| CD8a | 53-6.7 | PerCp Cy5.5 | 45-0081-82 | 300 | eBioscience |
| CD11b | M1/70 | PETxred/eF610 | 61-0112-82 | 300 | Invitrogen |
| Rat IgG1 | EBRG1 | FITC | 11-4301-82 | 300 | eBioscience |
| Rat IgG1 | EBRG1 | PE | 12-4301-82 | 100 | eBioscience |
| Rat IgG1 | EBRG1 | APC | 17-4301-82 | 200 | eBioscience |
| CD3e | 145-2C11 | eF610 | 61-0031-82 | 200 | eBioscience |
| Cd5 | 53-7.3 | PE Dazzle 594 | 100644 | 200 | Biolegend |
| Tbet | eBio4B10 | ef660 | 50-5825-82 | 200 | eBioscience |
| Foxp3 | FJK-16s | ef450 | 48-5773-82 | 200 | eBioscience |
| Rorgt | B2D | PE | 12-6981-82 | 75 | eBioscience |
| CTLA4 | UC10-4B9 | PE Dazzle 594 | 106317 | 100 | Biolegend |
| ICOS | 15F9 | PE | 107706 | 100 | Biolegend |
| PD-1 | J43 | PE Cy7 | 25-9985-82 | 200 | eBioscience |
| CD25 | PC61.5 | PE Cy7 | 25-0251-82 | 100 | eBioscience |
| Cx3cr1 | SA011F11 | PE | 149006 | 100 | Biolegend |
| CD64 | X54-5/7.1 | BV650 | 740622 | 100 | BD Biosciences |
| EpCam | G8.8 | eF450 | 48-5791-82 | 300 | eBioscience |
| iNOS | CXNFT | APC | 17-5920-82 | 200 | eBioscience |
| MHC-II | AF6-120.1 | PE | 12-5320-82 | 400 | eBioscience |

**Supplementary Table 3:** Antibodies used for flow cytometry experiments.
