## Supplementary figures and images for "Immunotherapy restores therapeutic efficacy of fecal microbiome transplants to treat *Clostridioides difficile*"

### Supplemental Figures

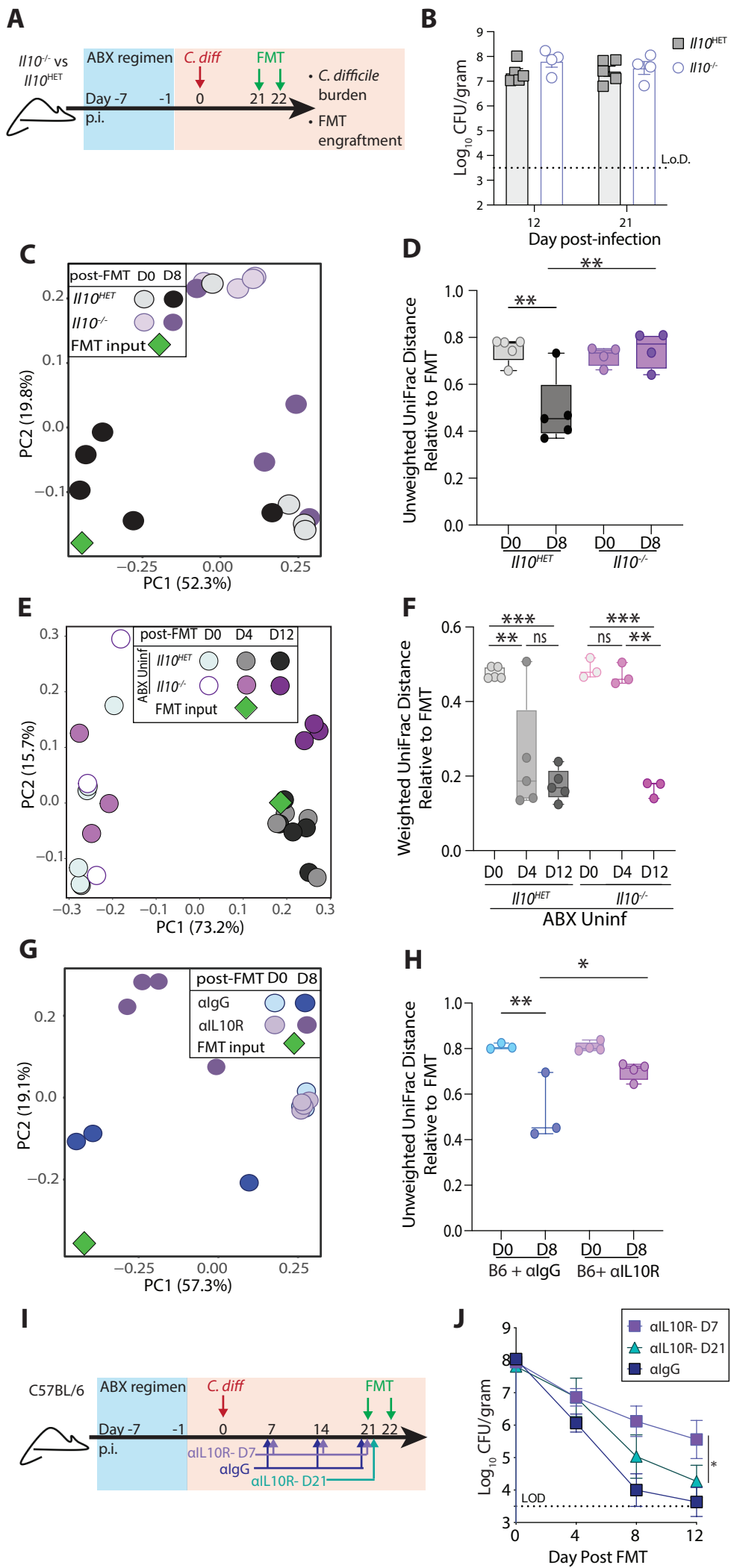

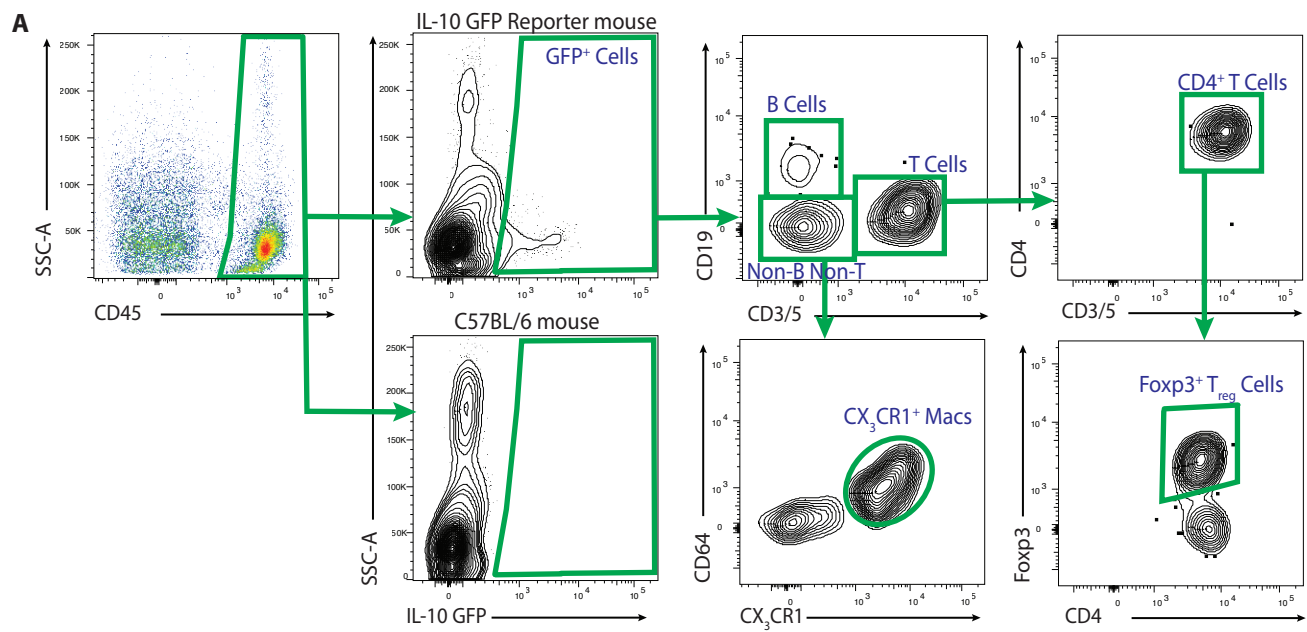

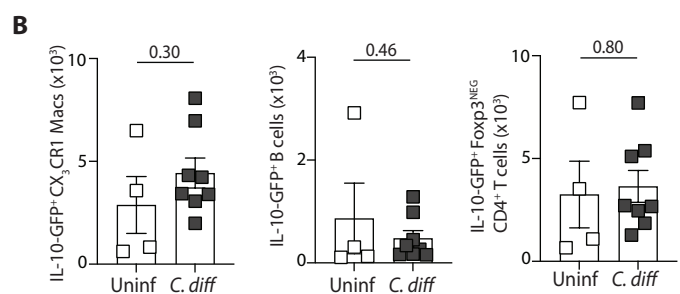

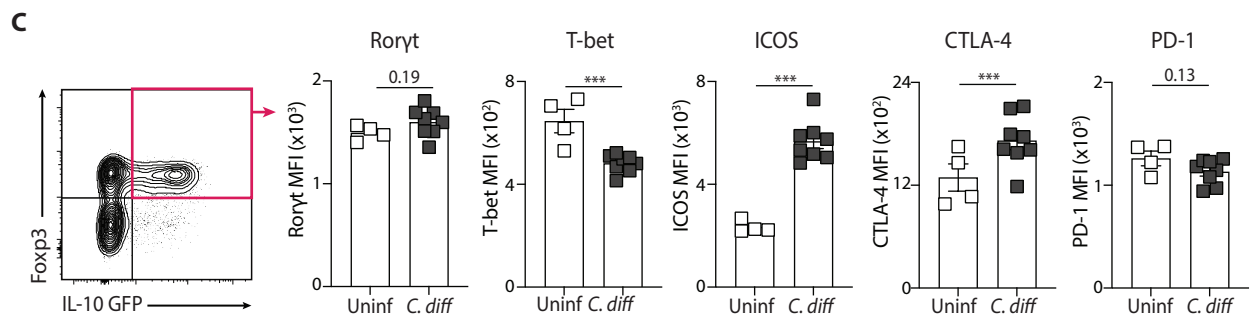

**A**

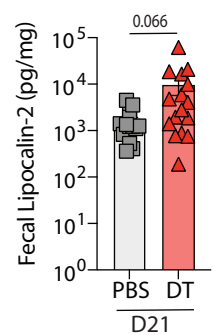

**B**

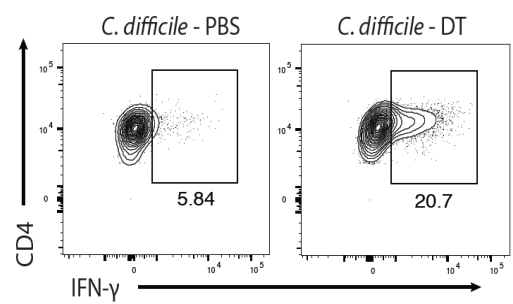

**C**

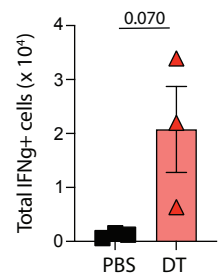

**D**

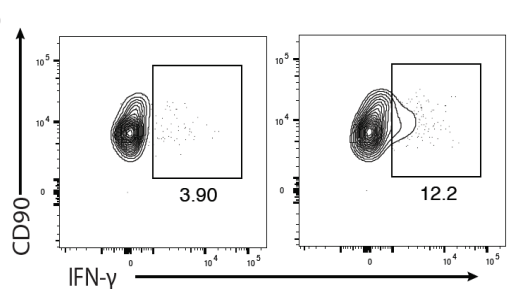

**E**

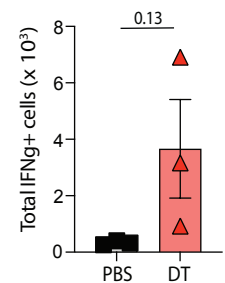

**F**

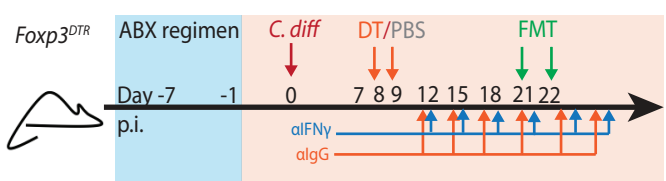

**G**

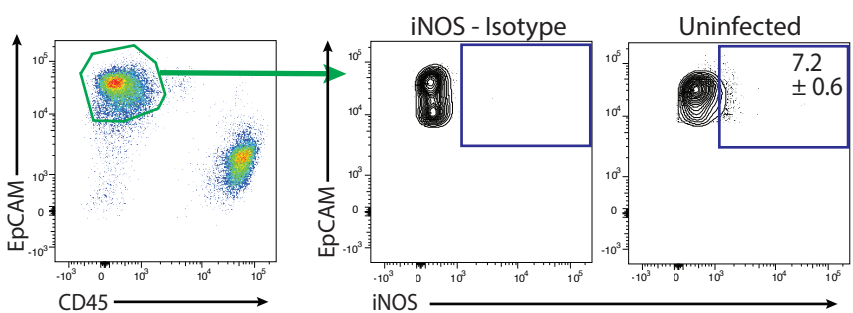

**A**

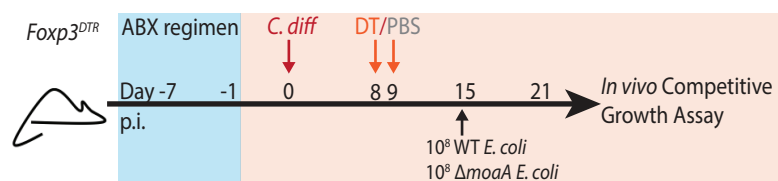

**B**

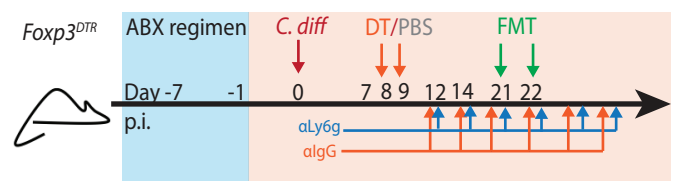

**C**

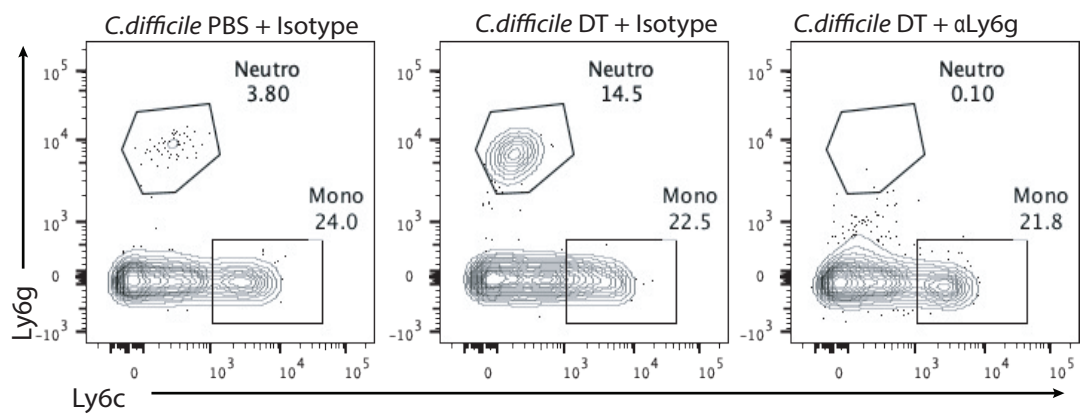

**D**

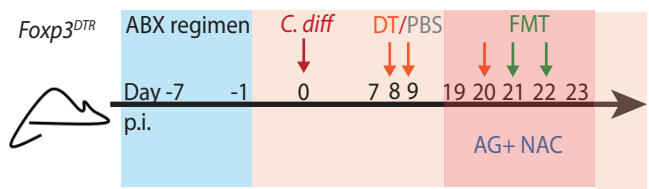

**E**

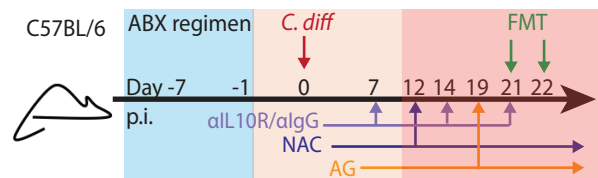

**A**

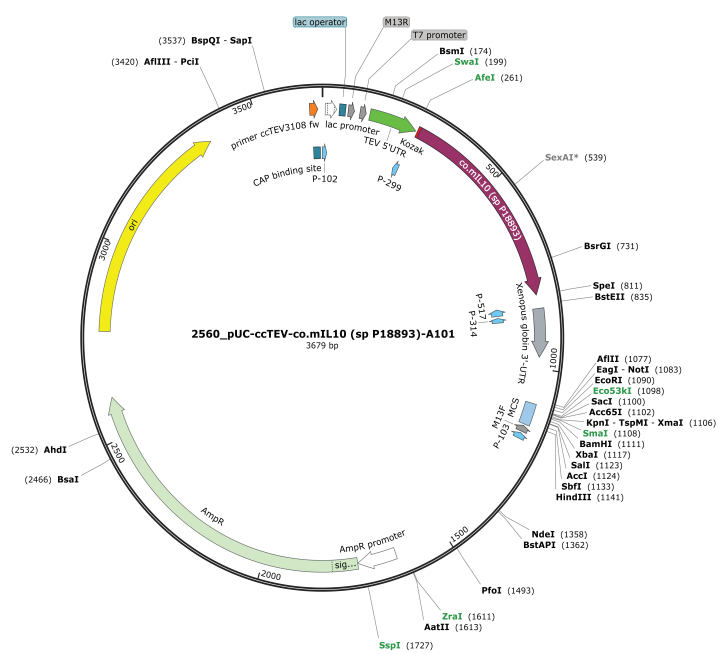

**B**

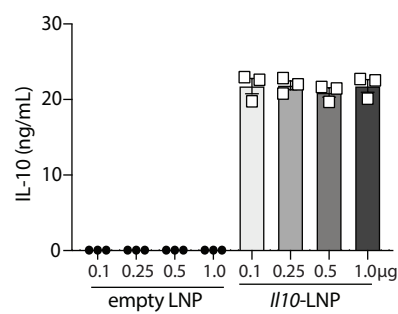

**C**

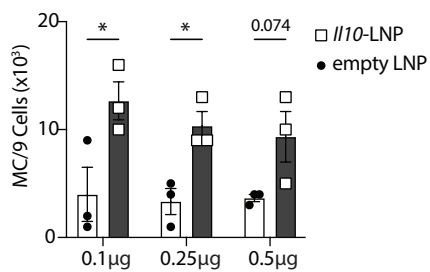

**D**

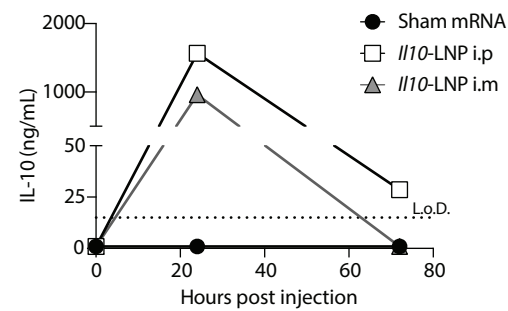

**E**

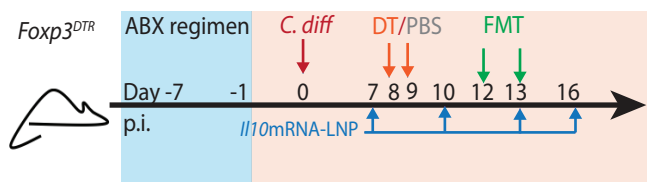

**F**

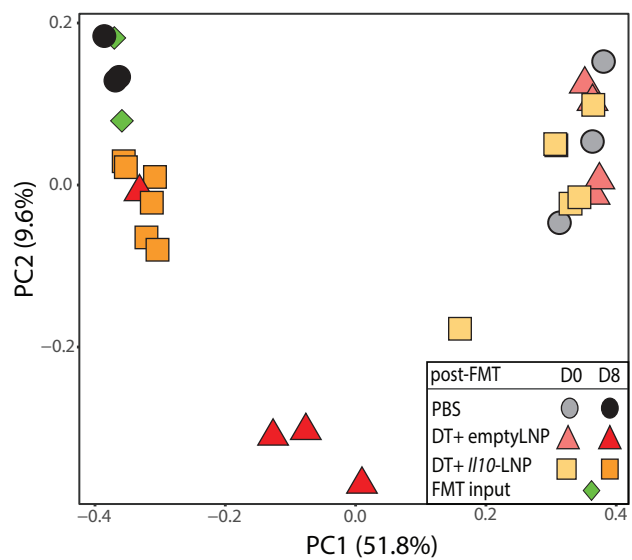

**G**

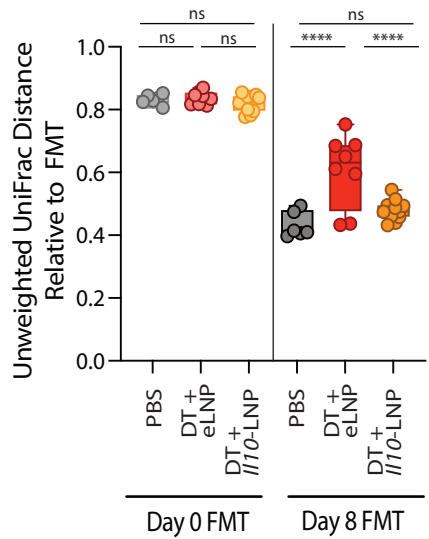
